# Addressing multiple facets of ligand-receptor network inference including single-cell proteomics

**DOI:** 10.1101/2025.10.05.680519

**Authors:** Jean-Philippe Villemin, Pierre Giroux, Morgan Maillard, Pierre-Emmanuel Colombo, Christel Larbouret, Jacques Colinge

## Abstract

Distinct ligand-receptor interaction (LRI) inference tools often produce markedly different results, and their performance can vary considerably across datasets. Indeed, performance is influenced by differences in experimental designs and dataset-specific features, making it difficult to establish a universal LRI tool. To address this challenge, we expanded our SingleCellSignalR Bioconductor package to provide an integrated framework that incorporates alternative scoring strategies and adjustable analytical depth. We motivate this choice through the analysis of two single-cell transcriptomics datasets that exemplify contrasting experimental designs. Leveraging the new framework flexibility, we present a detailed analysis of paired single-cell proteomics and transcriptomics data, providing, to our knowledge, the first direct comparison of LRI inference across these complementary modalities at single-cell resolution. Finally, we demonstrate how the same framework seamlessly accommodates additional underexplored data types from the LRI perspective, including patient-derived mouse xenografts and bulk RNA sequencing of upstream-sorted cell populations.

## INTRODUCTION

Mapping and exploring cellular networks mediated by ligand–receptor interactions (LRIs) is an important component of many single-cell transcriptomics (SCT) studies. To meet this need, a range of tools has been developed, each relying on a different strategy and sometimes addressing a different question [1,2]. Among the tools that infer LRIs from SCT data, we can distinguish two main categories: those based on gene co-expression or correlation, and those leveraging differential gene expression analysis. The first category tends to identify globally valid LRIs, while the second is tailored to identify enriched or exacerbated LRIs in specific cell populations [1]. A further distinction exists between tools that base inference on the ligand and receptor data only and those that also consider downstream target genes thought to be regulated by receptor activation [1,2]. Comparative benchmarks have consistently revealed limited overlap between different tool predictions [3,4]. Given the diversity of inference strategies, experimental designs and data modalities, it is unsurprising that a tool effective in one context may perform less well in another.

Although SCT technologies have revolutionized our ability to explore tissue composition, cell population heterogeneity, and cellular networks [5–9], limitations remain. Indeed, most biological processes are regulated by protein networks [10,11], but there are significant discrepancies between transcript and protein abundances [12,13]. Furthermore, transcriptomics does not bring any information on posttranslational modifications. These issues call for the development of single-cell proteomics as a complement or alternative to SCT. Few years ago, the Slavov group developed unbiased, MS-based single-cell proteomics (scProt-MS) [14]. With this technology, they could profile the expression of ∼1,000 proteins in 200 cells, which triggered many developments to address all sorts of technical challenges related to the use of very small amounts of material and to increase throughput [15,16]. The latest MS instrument developments now enable detecting deeper single-cell proteomes with ∼3,000 proteins [17–19]. As a result, although less mature than SCT, scProt-MS has reached a point where it can deliver useful and unique information [16–19,19–21] and it is time to assess its potential to unravel cellular networks. Nevertheless, due to their mode of acquisition, scProt-MS data harbor very different properties compared to SCT, thus adding to the data diversity we mentioned regarding SCT.

Moreover, other data types remain insufficiently addressed despite their practical relevance. For instance, FACS can isolate chosen cell populations for a particular study before analysis by bulk RNA-sequencing [22] or proteomics. Although cell-to-cell variability is averaged out, repeating such analysis across multiple samples results in count matrices conceptually close to SCT data that should be amenable to LRI analysis. However, such data also differ from SCT in fundamental ways exhibiting much higher dynamics, limited dropouts, and much lower replicate numbers. Another example is patient-derived xenografts (PDX). It is well-known that in PDX, after a few passages, the stromal cells are replaced by murine populations while patient human cancer cells remain. Dual alignment against human and mouse genomes thus enables segregation of cancer versus stromal transcriptomes [23], making it theoretically possible to investigate paracrine and autocrine LRIs across these compartments with SCT-like LRI algorithms.

Altogether, the question of inferring LRIs is hence not a single well-defined problem but rather a multifaceted challenge that calls for flexible strategies. Although using multiple tools or a federation of tools might provide a solution [3], it is complex to maintain. We preferred an integrated yet open strategy in which common data structures and functionalities support alternative LRI scoring schemes and data types. We first show the need for such flexibility through two illustrative SCT examples, where we also apply a selection of other tools. Since roughly 100 different LRI tools exist [2], and our purpose was not to provide a thorough benchmark, we selected a few recent tools considering intracellular signaling and either differential gene expression or co-expression. We then demonstrate the application of the new functionalities and differential model of SingleCellSignalR. Our main focus is the analysis of an scProt-MS dataset and its paired SCT data. Finally, we illustrate our framework ability to handle cell-sorted bulk transcriptomes and PDX data with equal ease.

## MATERIALS AND METHODS

### Reference databases

In its first version, SingleCellSignalR [9] relied on a compilation of KEGG [24] and Reactome [25] pathways to assemble a reference interactome. This compilation was provided by Pathway Commons [26]. The LRI database LR*db* was compiled by ourselves from multiple sources [9], and the definition of pathways as sets of genes or proteins was provided by Reactome pathways and Gene Ontology Biological Process (GOBP) terms [27]. To ensure updates at regular and controlled frequencies, we have introduced a different procedure to compile equivalent databases. The reference interactome is now generated by a Python script that directly interrogate a local installation of Reactome neo4j database. We further add gold-rated transcription factor (TF)-gene interactions from ExTRI [28] (Fig. 1A). The same Python script interrogating Reactome outputs putative LRIs. By using the original 2019 LR*db* as a basis, these LRIs are simply added provided new and following manual inspection (Fig. 1A). The gene sets defining pathways are still taken from Reactome and GOBP, which we now retrieve through MSigDB [29] to provide compatibility with popular gene set enrichment tools. In every case, *Homo sapiens* data were used only. BulkSignalR built-in mechanism to map orthologous genes of other species to *Homo sapiens* must be used for nonhuman data.

**Figure 1.**
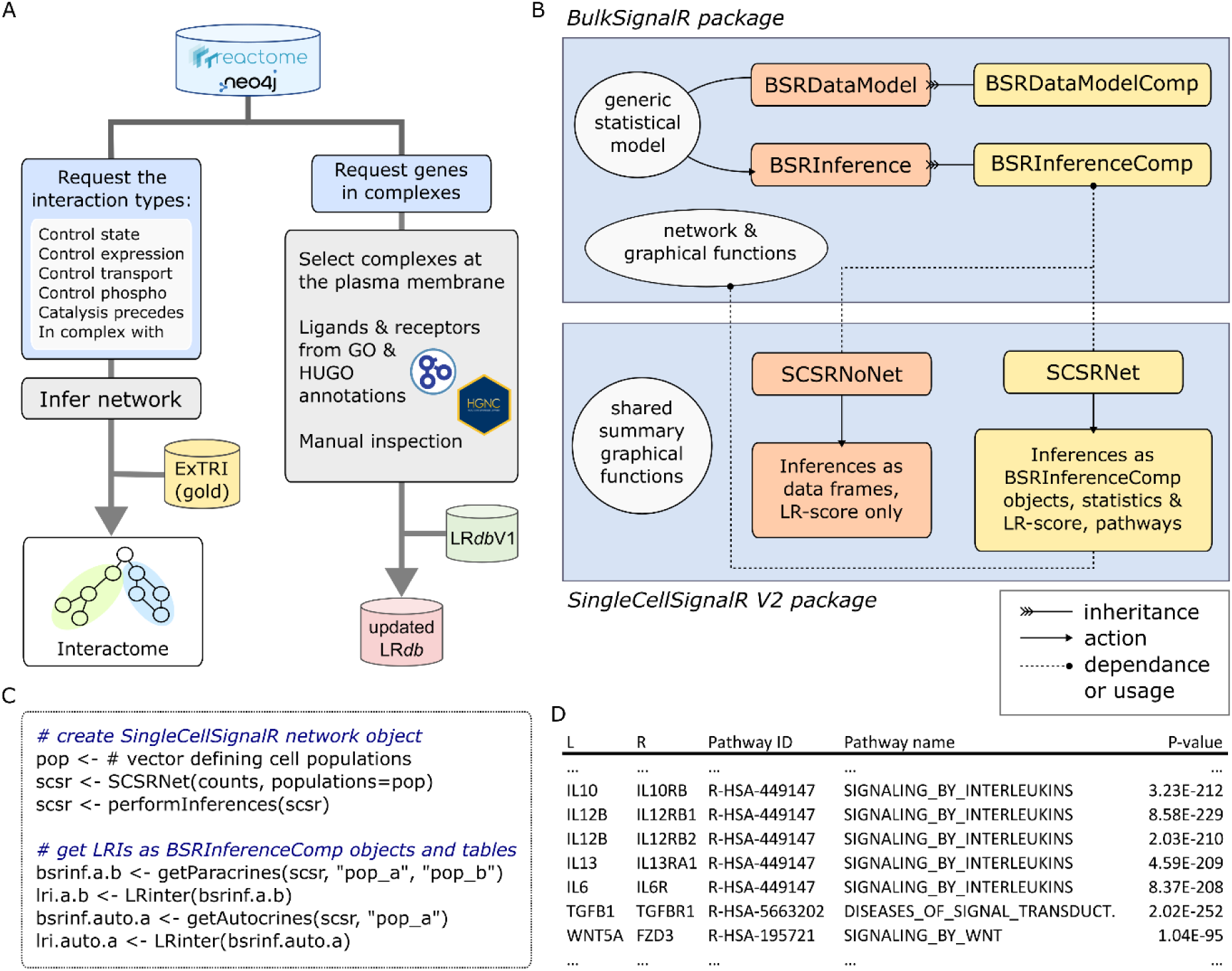
SingleCellSignalR V2 design. (**A**) Reference interactome generation and LR*db* update process overview. (**B**) High-level dependencies between BulkSignalR and SingleCellSignalR S4 classes. (**C**) Example R code to create the basic objects and generate inferences. (**D**) Illustrative inference (part) after conversion into a table with the accessor LRinter() as in (C).

Reactome defines protein complexes and it also covers a range of metabolites that are involved in diverse reactions. While proteins in a complex are all connected to each other, this natural strategy becomes problematic for a few gigantic complexes in Reactome that potentially cause excessive connectivity. We thus set a threshold on complex sizes, beyond which they are ignored (Fig. S1A). The current threshold is at 10. Similarly, metabolites participating in very large numbers of interactions such as ATP or water induce connections between functionally unrelated genes. We thus also set a threshold on metabolites based on the number of interactions they participate to (Fig. S1B). The current threshold is at 50. Provided reasonable values were imposed, we have not observed major variations in terms of which LRI were detected. Therefore, we only applied one level of filtering with the aforementioned thresholds.

Obviously, the different sources are updated before we launch our own update. In its version 2, SingleCellSignalR accesses the reference databases from BulkSignalR [30]. The latter package downloads the databases from a FTP server we set up for this purpose. Upon first installation, BulkSignalR obtains the latest database versions (2025 at the time of writing). In subsequent uses, it checks whether a new version is available and issues a message if this is the case. The user can then decide to trigger the update, or to stay with the originally downloaded version. In every case, users can download previous versions of the databases from our FTP server or substitute their own, provided they respect our simple tab-based formats and interaction ontology. Users can also add chosen LRIs that would be missing, or which they want to challenge with gene or protein expression data. The latest 2025 version of the databases included 761 ligands, 702 receptors for 3,361 known LIRs. The 2024 version, which was used to prepare most examples and figures, has its parameters reported in Table S1.

### Software design

Version 2 of SingleCellSignalR builds upon its bulk and medium-resolution spatial data counterpart BulkSignalR [30]. Indeed, a key new feature of SingleCellSignalR version 2 is the ability to incorporate biological pathways downstream receptors for more accurate statistical modeling. Since this feature was central to BulkSignalR, we implemented SingleCellSignalR version 2 as a software layer on top of it. This design allows SingleCellSignalR version 2 to perform integrated statistical modeling of (ligand, receptor, downstream pathway) triples (see Results for details). In parallel, SingleCellSignalR original scoring called the LR-score is still computed for compatibility, and also because there are contexts where it outperforms statistical modeling (see Results). Both BulkSignalR and SingleCellSignalR are included in Bioconductor and share an object-oriented design based on S4 classes. An overview of the relationship between the two packages, their main classes, and functionalities is provided in Figure 1B.

BulkSignalR BSRDataModel is a container for expression data, including transcriptomics or proteomics in the bulk, single-cell, or medium-resolution spatial formats. Through the application of a generic statistical model capturing the significance of triples (ligand, receptor, downstream pathway target genes), LRIs are inferred that are represented in a BSRInference object. The inference relies on correlation analysis and is meant for bulk or medium-resolution spatial expression data such as 10x Genomics^TM^ Visium (see BulkSignalR publication [30] for details). The daughter classes BSRDataModelComp and BSRInferenceComp enable the inference of LRIs through differential analyses between user-defined sample groups or clusters. This alternative approach allows the identification of LRIs that are enriched in specific clusters, in contrast to globally inferred LRIs through correlation analysis over all the samples. Both inference modes rely on the same statistical model, which only needs the significance (P-values) of observations (on ligands, receptors, and target genes in pathways) [30].

Single-cell functionalities are implemented through the SCSRNet class, which automatically generates both paracrine LRI inferences between cell population and autocrine LRI inferences within individual populations. SCSRNet inferences are based on BulkSignalR sample cluster comparisons. By default for each population, gene or protein expression is contrasted against all the other populations in the dataset to find regulated ligands, receptors, and receptor targets. Autocrine LRIs can be inferred from one such differential analysis with actual computations carried out by BSRInferenceComp objects. By contrast, paracrine LRI inferences require two differential analyses: one at the source population to identify regulated ligands, and another one at the target population to identify regulated receptors and receptor target genes in downstream pathways. This is also obtained through BSRInferenceComp objects, which can accommodate paired source and target population differential analyses to support paracrine inference. Importantly, users may substitute the default differential analysis with custom logic and/or statistical tests tailored to specific experimental designs or data types. Default differential analysis is detailed under statistical model below. Since SCSRNet-derived LRI inferences are returned as BSRInferenceComp objects, all visualization and network extraction functions available in BulkSignalR can be directly exploited.

As a complement to SCSRNet, we implemented a no-network variant named SCSRNoNet, which generates paracrine and autocrine LRIs without incorporating receptor downstream signaling (Figs 1C-D). This class only relies on the LR-score introduced in SingleCellSignalR version 1, whereas SCSRNet does both the statistical modeling with pathways and LR-score computations. The SCSRNoNet class allows reproducing SingleCellSignalR version 1 results in this new implementation, and it is much faster than SCSRNet. SCSRNoNet LRIs are returned as tables (data frames) directly contrary to SCSRNet that stores LRIs in BSRInferenceComp dedicated data structures and requires an accessor, LRinter(), see Figure 1C, to obtain tabular summaries.

### Statistical model

The statistical model is detailed in BulkSignalR paper [30]. We briefly recall its principle and explain how it was adapted to differential expression analysis. In BulkSignalR correlation analyses, null distributions are directly learned from data using a random permutation scheme: a density function 𝑓_𝐿𝑅_: [−1; 1] → ℝ^+^ for the ligand-receptor random correlations, and a density function 𝑓_𝑅𝑇_: [−1; 1] → ℝ^+^ for the receptor-gene target random correlations. Ligand-receptor correlation significance is given by 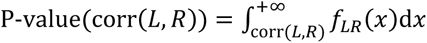. Similarly, for a specific gene target 𝑇, significance of its correlation with the receptor is given by 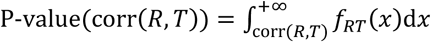. To combine the correlations between the receptor and each individual gene target in a specific pathway 𝑊, we consider the number of targets 𝑁 and the rank we want to consider. The rational is that using the best correlation or the worst might not reflect well actual relationship, and to try to model each pathway in a sophisticated manner might not be robust given missing data, diversity in pathway types, and errors or absence from the pathway definitions. By using something such as the median correlation, or the median correlation P-value actually, we obtain a robust and simple schema. To be more flexible, the rank of the retained P-value is a free parameter of the model that is defined as a proportion of the pathway size 𝑁. By default it is 0.55 × 𝑁. The P-value of this rank 𝑁 observation is obtained by using rank statistics, thereby making the simplifying assumption that all the receptor-target correlations are independent observations under the null hypothesis. We denote this rank P-value as P-value(𝑊), see BulkSignalR paper [30] for details. Altogether, the correlation-based P-value of a (ligand, receptor, pathway) triple is provided by P-value(corr(𝑅, 𝑇)) × P-value(𝑊), assuming independence of the two observations under the null hypothesis.

In case differential expression is chosen, then a P-value representing the significance of regulation of each involved molecule is obtained from the differential analysis. P-values of the gene targets in 𝑊 are integrated as above using a rank statistics, and the overall P-value is provided by P-value(𝐿) × P-value(𝑅) × P-value(𝑊), where similar simplifying assumptions are made under the null hypothesis. Note that is this case, functions equivalent to 𝑓_𝐿𝑅_ and 𝑓_𝑅𝑇_ are no longer necessary. The statistical model behind the differential analysis can be chosen by the user. By default, we use a simple 2-sidded Wilcoxon test that is rather common in SCT. The direct application of Wilcoxon will of course depend on the sizes of the cell populations we compare, which is misleading in the context of a tool that should treat each population with comparable sensitivity. To circumvent this issue, our default procedure resamples 10 times each population (with replacement) and perform Wilcoxon 50 cells *versus* 50 cells. The results (P-values, fold-changes) are averaged to obtain a final analysis that is cell population size independent. Parameters of this resampling procedure can be adjusted by the user.

### Other software packages and reference databases

Whenever possible, we used the default parameters of each algorithm and tried to run them with identical or as close as possible reference databases. We ran cellChat [31] using the CellChatdb.human database and set the type parameter of the computeCommunProb() function to trimean. To execute scSeqComm [32], we selected the scSeqComm package-provided LR_pairs_CabelloAguilar_2020 file as the reference LRI database, TF_TG_TRRUSTv2_HTRIdb_RegNetwork_High as the TF to gene targets database and TF_PPR_REACTOME_human as the reference for receptors-TF associations. To launch scMLnet [33], we used its default reference files. We called CytoTalk [34] using our own version of LR*db* (2024 version) and HUGO gene symbols as reference. SingleCellSignalR version 2 was used with the 2024 version of our databases.

### CD40L-CD40 dataset

This dataset [35] was available from Gene Expression Omnibus (GEO) database [36] under the reference GSE228415. We kept features detected in at least 3 cells and included cells harboring at least 200 features using parameters of the CreateSeuratObject function from Seurat R package [37]. Cells with fewer than 750 or more than 15,000 unique molecular identifiers (UMIs), or with >20% of reads mapping to mitochondrial or ribosomal genes, were excluded. Putative multiplets were filtered out by removing cells with >75 detected genes *per* 100 UMIs. We were left with 5,018, 4,645, and 6,001 cells for GFP_GFP, CD40L_GFP, CD40L_CD40 experiments, respectively. We merged the data slots and joined the layers of the different Seurat objects (merge/JoinLayers) and UMI counts were normalized with SCTransform, except for scMLnet, which requires raw counts. Natural and exogenous CD40LG and CD40 levels were summed separately. We next annotated cells as B cells, monocyte, natural killer (NK) cells, CD4+ T cells, and CD8+ T cells using singleR [38] and an annotated reference of immune cells from celldex Bioconductor package [38] from the same authors. The number of cells in each population for each condition is reported in Table S2.

During data analysis, cellChat LRI significance was defined with P-value < 0.05. We selected scSeqComm LRIs with S_inter > 0.9. ScMLnet LRIs were imposed P-values < 0.05 and a log-fold-change (logFC) > 0.15. No threshold must be set for CytoTalk that returns LRIs based on comparisons with random networks. We used it as such. SCSRNoNet LR-score threshold was set to 0.85. SCSRNet LRIs were imposed a maximum ligand and receptor P-value = 0.01, a minimum absolute logFC = log_2_(1.01), with pathway gene targets minimum positive logFC = log_2_(1.01). Note that on purpose SCSRNoNet was imposed a rather strict threshold [9] and essentially no selection was imposed to SCSRNet.

### Lung cell atlases

From the integrated Human Lung Cell Atlas (HLCA) [39], we two datasets: Barbry_Leroy_2020 and Nawijn_2021, both generated with 10x Genomics Visium technology. We used the original cell annotations and retained the healthy, respiratory airways cell populations with at least 50 cells in both atlases and marked as correctly annotated. See Table S3 for the respective population sizes. Raw counts were normalized using Seurat SCtransform function from Seurat, but for scMLnet that needs raw counts. The various tools used for comparisons were used with parameters as in the CD40-CD40L dataset, except for SCSRNet, where ligand, receptor and target gene minimum logFC was set to log2(1.1).

### MONO-MACRO scProt-MS and SCT datasets

These datasets were released by Specht, et al. [18]. ScProt-MS data were downloaded from the Slavov’s group website directly as indicated in the original publication. They are also available from MassIVE (IDs MSV000083945 and MSV000084660). Mapping UniprotKB/Swissprot IDs to HUGO gene symbols was based on the Swissprot version downloaded on March 30, 2023. Hierarchical clustering was done with R hclust function using the ward.D method and the Euclidean distance. The 2019 version of our reference databases was used for this particular dataset. SCT data (two replicates) were downloaded from GEO (GSM4226877 and GSM4226878). Genes with UMI counts > 0 were considered expressed, and cells with 3,500 to 6,000 expressed genes were selected (Figure S2A).

Genes with UMI > 1 in at least 15% of the selected cells were considered, and the other genes were discarded. This left us with 4,296 genes in 7,177 cells. No batch correction for the replicates was applied because no batch effect was visible (Fig. S2B-C).

Both scProt-MS and scRNA-seq data were analyzed with SingleCellSignalR. We substituted the default differential analysis with a simple function that was adapted to the data (cf. Fig. 1C, SCSRNet() constructor called with a user-defined function, see Supplemental Material). In the scProt-MS case, Wilcoxon tests were used for differential expression and log-fold-change were computed as differences to accommodate the particular nature of the data. In the scRNA-seq, we also defined a specific function given the particular experimental design, but it follows the default function logic otherwise. Resampling of data was made with a number of cells equal to scProt-MS (instead of 50) to make results more comparable.

### FACS-sorted bulk transcriptomics

This dataset was comprised of RNA sequencing data from mouse lung tumors and WT control animals where selected cell populations were FACS sorted before sequencing [22]. We downloaded the Fastq files from NIH SRA repository based on references in GEO GSE59831 entry. Sequences were aligned against *Mus musculus* genome (GRCm39) with STAR (version 2.7.1a), which also extracted gene read counts.

### PDX bulk transcriptomics data

Generation of the PDX and transcriptomic data is detailed in our recent papers [40,41], including patient-derived cell lines, animal work, sequencing and genome alignment. Relevant ethical agreements were obtained. We do not repeat these details here. We also used one control sample made of healthy human pancreas (h_hPanc) obtained from ICM clinical-biological bank (numbers (PROICM2018-10 BPA; ID RCB 2018-A02901-54). We also used as murine control sample brown adipose tissue (m_BAT). RNA-sequencing data were submitted to GEO.

## RESULTS

### Comparing related cell populations

A number of SCT investigations aim at exploring the heterogeneity of a single broad cell population or of closely related populations such as cancer-associated fibroblast [42,43] in tumors, or subsets of immune cells in autoimmune diseases. Such experimental designs usually involve cell sorting prior to SCT, resulting in datasets with limited transcriptional profile diversity. This poses a challenge for LRI inference tools that rely on differential gene expression.

To illustrate this, we analyzed a dataset in which a ligand (CD40L) and its cognate receptor (CD40) expression were artificially induced in specific cell populations [35]. Natural killer (NK) cells – normally lacking CD40L expression – were transfected with mRNA encoding CD40L, while B cells – normally expressing only low levels of CD40 – were transfected with mRNA encoding the receptor. The dataset further included monocytes and T cells, and multiple experimental batches were produced covering all possible combinations of mRNA transfection, with GFP mRNA serving as a control. We applied the different inference tools to the whole dataset and restricted our analysis to NK and B cell populations in the CD40L- and CD40-transfected conditions. The expected CD40L-CD40 interaction was only detected by tools relying on transcript expression without differential analysis, namely cellChat, CytoTalk, and SCSRNoNet. In contrast, differential analysis-based approaches, *i.e.*, SCSRNet, scMLNet, and scSeqComm, failed to recover this interaction (Fig. 2A). Thresholds imposed to each tool to select LRIs are reported in Materials and Methods. Of note, SCSRNet was essentially imposed no selection, while selection was quite strict for SCSRNoNet LR-score to ensure results for our framework were not obtained by chance. Also note that the different tools predicted varying numbers of LRIs. In the absence of ground truth, we cannot interpret these counts.

**Figure 2.**
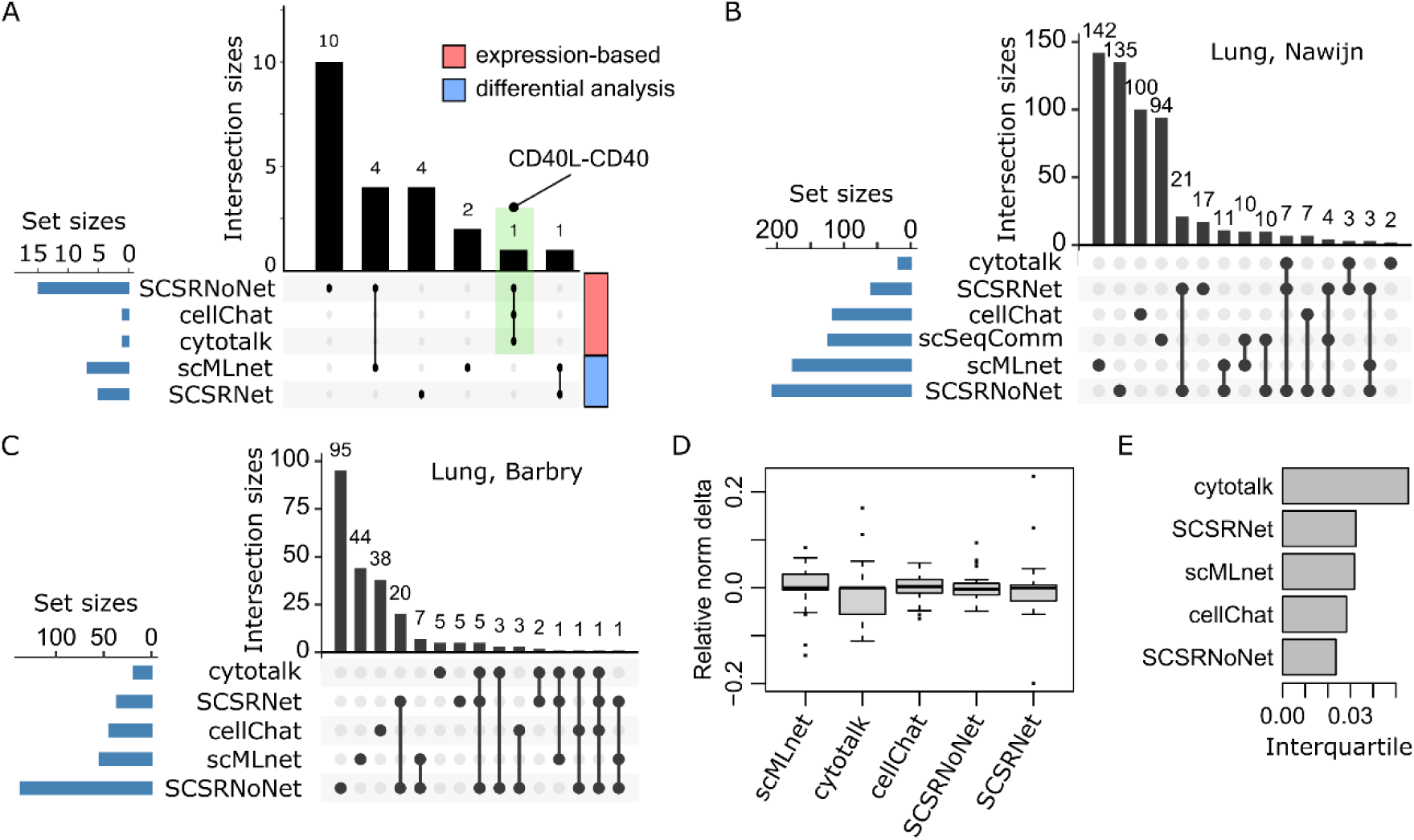
SCT comparisons. (**A**) UpSet plot featuring the number of detected LRIs from the CD40L-CD40 dataset. The scSeqComm tool did not identify any significant LRI, it is thus absent from the plot. SCSRNoNet stands for SingleCellSignalR LR-score, SCSRNet stands for SingleCellSignalR with pathways, differential analysis and statistical modeling. (**B**) UpSet plot for Nawijn, *et al.*, lung atlas considering four main cell populations. (**C**) Same for Barbry, *et al.*, lung atlas. (**D**) Relative normalize differences between the number of LRIs inferred for the same cell populations in the two atlases (numbers normalized to Nawijn data, normalized differences divided by the total LRI inferences of each tool). (**E**) Interquartile difference for each tool in (**D**) boxplot.

### Comparing diverse cell populations

To represent experiments harboring diversity in the sequenced cell populations, we decided to use a single-cell atlas for a chosen tissue. More specifically, we also wanted to assess the question of result robustness when similar data for the same tissue were analyzed. Therefore, we selected the two largest (after correctly annotated filter) respiratory airway cell atlases from HLCA [39]. To facilitate the robustness study, we limited ourselves to populations present in both atlases with at least 50 annotated cells. Namely, we considered paracrine LRIs between B cells, plasma cells, respiratory basal cells, mast cells, mucus secreting cells, multiciliate epithelial cells. Table S3 reports the number of cells in each atlas selection. Figures 2BC show the number of LRIs identified by each tool as well as the overlap between tools in the two datasets. There was clear discrepancy between the various tools as it is essentially always the case with LRI inference. Interestingly, although the datasets were comparable, the total number of LRIs found by each tool also varied substantially. An extreme case was scSeqComm, which did not return any LRI with Barbry, *et al.* data. To characterize robustness further, we compared the LRIs identified in each of the 30 possible pairs of cell populations in each dataset. Adjusting for the total *per* tool by a factor to make the numbers in each dataset comparable, we computed the difference. Because each tool retrieved a different total number of LRIs, we divided the normalized differences by the tool total (Fig. 2D). In this analysis, the smallest variation was displayed by SingleCellSignalR LR-score (SCSRNoNet, Fig. 2E), which was also the scoring identifying the largest number of LRIs. Conversely, CytoTalk, which identified the smallest number of LRIs, was the less robust. See also Figure S3 for absolute differences.

### ScProt-MS data and their specificities

To understand the nature of the data generated using scProt-MS, we provide a brief overview of the main used workflows. Classically, scProt-MS protocols require to dissociate cells when starting with a solid tissue (Fig. 3A). Then, cells are isolated and each cell is deposited in one well of a microtiter plate for the preparation steps, such as lysis, denaturation, and digestion, before the proteomic analysis. Then, two main routes are available that diverge from SCT: i) small groups of 𝑛 cells are pooled together for multiplex MS analysis by taking advantage of isobaric tags, *e.g.*, 16-plex tandem mass tags (TMT), or ii) each single cell is analyzed by MS.

**Figure 3.**
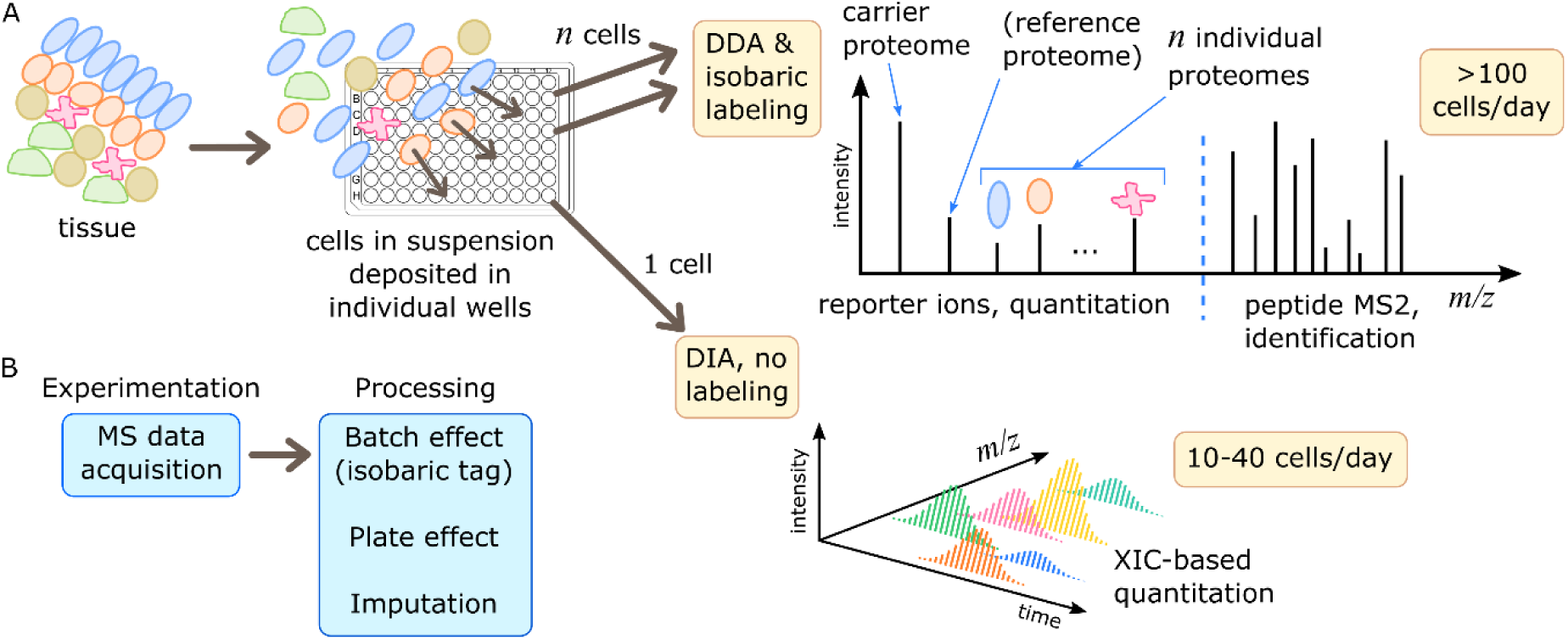
Single-cell proteomics workflows. (**A**) Typically, cells in a solid tissue are dissociated by enzymatic digestion (not needed for circulating cells) before deposition in the wells of a microtiter plate. In the plate, cells are prepared individually (e.g. lysis, digestion, labeling) for the MS analysis. Currently, there are two classical, generic workflows: i) pools of cells are analyzed simultaneously using an isobaric tag system and data-dependent acquisition (DDA) MS, and ii) cells are analyzed individually in data-independent acquisition (DIA) MS, where quantitation relies on the areas under the extracted-ion chromatogram (XIC). (**B**) After acquisition, data are normalized to compensate for various biases and imputation is carried out to replace missing values.

In the first typical workflow, named SCoPE-MS [14], a so-called carrier proteome at a concentration equivalent to 100-200 individual cells, is assigned to a specific channel of the isobaric tag system. The carrier proteome composition should represent the average of all cell types in the tissue under study. Its role is to boost peptide detection by increasing MS signals (Fig. 3A). In some cases, another channel is assigned to a reference proteome used to normalize data between pools of 𝑛 cells. The remaining isobaric tag channels are used for individual cells. MS is carried out in data-dependent acquisition (DDA) mode. Besides augmenting MS signals thanks to the carrier proteome, this approach increases throughput because the proteomes of several individual cells are acquired simultaneously [14,18,44]. Data require extensive normalization to correct the batch effects introduced by cell pooling and microtiter plates (Fig. 3B). Moreover, the number of missing values is high due to signal intensity differences between the carrier proteome channel and the individual cell channels and the generally low concentrations of proteins. These missing values are usually compensated by imputation. Overall, the final data resemble z-scores and low expression leads to negative values (Fig. S4), a situation very different from SCT data.

In the second typical workflow, individual cells are analyzed separately following a data-independent acquisition (DIA) MS strategy [45]. Unlike data-dependent acquisition (DDA), DIA does not select individual peptides for subsequent fragmentation, but fragments all precursor ions entering the mass spectrometer at a given time. It requires the prior preparation of a reference library (in DDA mode) of reporter MS2 peaks to identify the peptides. This reference library is generated from a large pool of cells that include all the populations present in the tissue under study at balanced concentrations. Usually, DIA can identify low-abundance peptides because of the absence of peptide selection that characterizes DDA. DIA-based scProt-MS data also require substantial imputation of missing values. Moreover, rather strong normalization is necessary to correct the individual sample variability and microplate effect (Fig. 3B). Qualitatively, DIA and DDA-isobaric tag scProt-MS data are hence not so different (Fig. S4).

Few authors proposed workflows that depart from these two main models, for instance, an integrated microfluidic device that performs cell isolation and DIA MS [46], and individual cell preparation in droplets instead of microtiter plates [47]. This usually does not change the nature of the obtained data.

### Proteome depth in scProt-MS data

The majority of the first scProt-MS datasets reached a proteome coverage of 1,000 to 1,300 proteins. While ligands and receptors are usually included in such shallow proteomes, it is seldom that members of a putative LRI are concomitantly identified, not to mention a sufficient number of targets under the receptor. Accordingly, at this depth, scProt-MS has limitation in inferring LRIs.

Nevertheless, in such datasets, we showed that substantial information about cell identity was specifically carried by ligands and receptors (Suppl. Results, Figs. S5-S6). Most recently, the introduction of the last generation of MS instruments allowed several teams to reach 2,000-3,500 proteins deep proteomes. At such a depth, tools such as SingleCellSignalR can be applied.

### A scProt-MS macrophage-monocyte dataset

The dataset authors incubated U-937 monocytes with phorbol-12-myristate-13-acetate (PMA), a protein kinase C agonist, to induce their differentiation into macrophage-like cells [18]. Untreated U-937 cells and differentiated macrophages were collected and analyzed using the SCoPE2 workflow that achieved a depth of 3,042 proteins in 1,490 individual cells. Mapping protein identifiers to HUGO gene symbols left us with 3,032 molecules, including 122 ligands and 89 receptors. We clustered the individual proteomes and compared this clustering with the cell type annotations provided by the authors (Fig. 4A). Untreated U-937 monocytes displayed a specific profile, whereas PMA-treated cells formed a gradient. We sorted cells in function vimentin (VIM) expression, which is associated with macrophage biology, annexin A2 (ANXA2), and suppressor of glucose, autophagy-associated protein 1 (SOGA1) [18]. This accurately separated untreated U-937 cells from macrophages (Fig. 4B). We empirically defined the 20% of cells at the bottom and top of vimentin expression as monocytes and macrophages, respectively.

**Figure 4.**
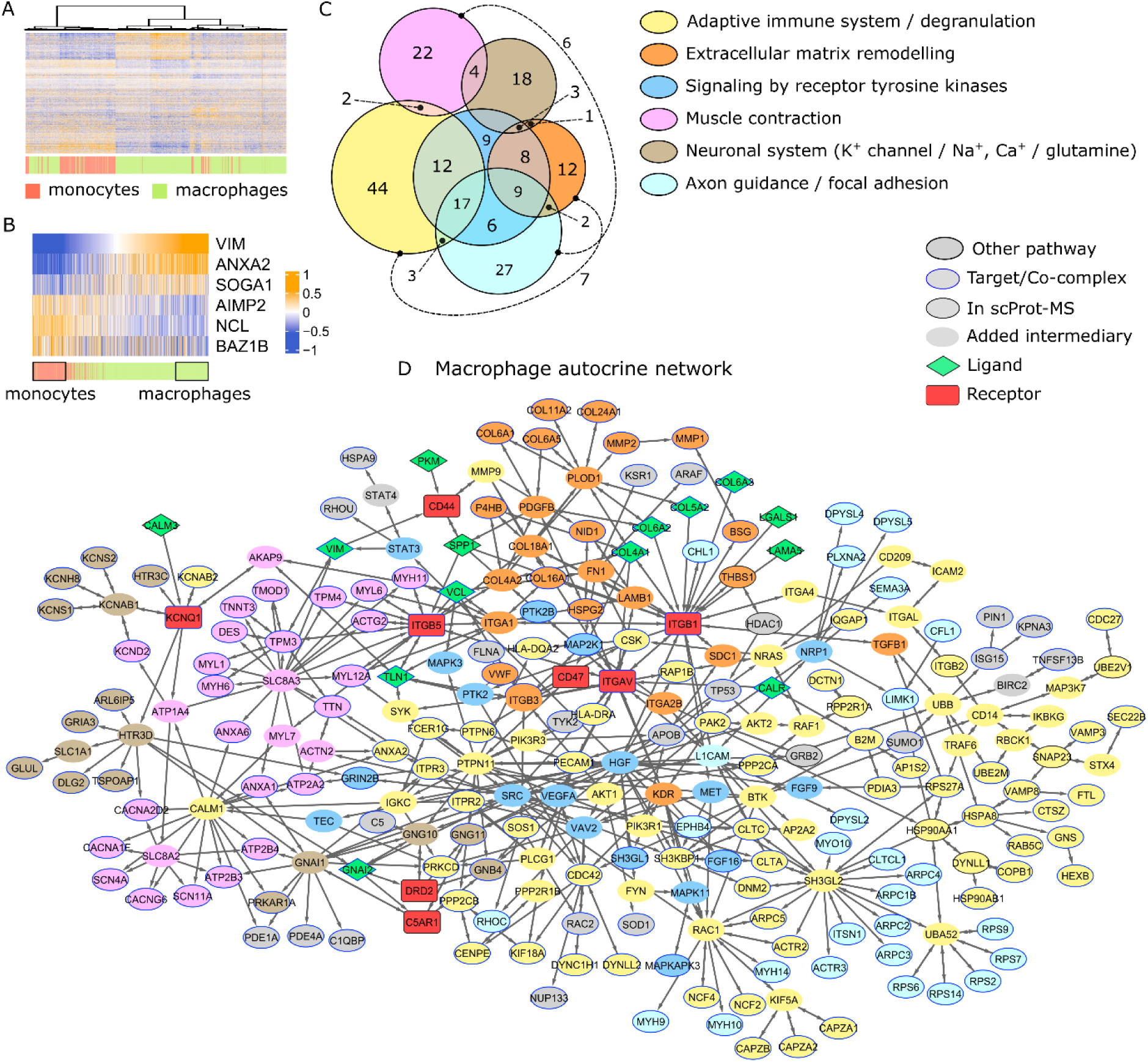
Autocrine ligand-receptor interactions in macrophages. (**A**) Cluster analysis of all the individual proteomes. Monocyte/macrophage annotations provided by the authors [18]. (**B**) Sorting in function of vimentin (VIM) expression allowed defining the top/bottom 20% of cells as macrophages/monocytes, respectively (black rectangles). (**C**) Main receptor downstream pathways with numbers of non-ligand and non-receptor proteins. The immunity and extracellular matrix (ECM) pathways shared eight proteins that could not be represented in the Venn diagram. See Figure S7 for the complete UpSet plot. (**D**) The LRI and downstream protein networks in macrophages. For proteins present in several pathways, color annotation priority was given to immunity and ECM, hence the limited number of receptor tyrosine kinase signaling-associated proteins (blue). The BulkSignalR algorithm links downstream target genes in pathways to the receptors through their shortest path. Some pathway proteins not detected can be added by this process, and they are featured without a border. All scProt-MS-detected proteins have a border. Target genes have a blue border. Note that some ligands or receptors were present in the identified pathways, but they were not implicated in any LRI. Such proteins are featured with the standard node shape, for instance ITGB2 or B2M.

We applied SingleCellSignalR differential approach to compare the macrophage and monocyte proteomes with the aim of identifying autocrine LRIs and the associated pathways upregulated in macrophages. Differential expression analysis was adapted to the data (Materials and Methods). We found 14 ligands and 8 receptors with increased expression in macrophages that formed 22 significant LRIs linked to regulated targets in pathways downstream the receptor according to our statistical model (Supplementary Data). Of note, among the other tools we tested above, only cellChat could handle scProt-MS data returning 42 LRIs. Since cellChat is not based on differential analysis, *i.e.*, enriched pathways, we did not analyze its output further.

We next exploited BulkSignalR network functionality to generate a parsimonious molecular network that linked receptors to their targets through the shortest paths in the identified pathways. Eight predominant pathways covered most of the proteins in this network: adaptive immune system and neutrophil degranulation (Reactome IDs R-HSA-1280218 and R-HSA-6798695), extracellular matrix (ECM) remodeling (R-HSA-1474244, R-HSA-3000178, R-HSA-1474228, R-HSA-3000171), muscle contraction (R-HSA-397014, R-HSA-445355), signaling by receptor tyrosine kinase (RTK) (R-HSA-9006934), neuronal system (R-HSA-112316), axon guidance (R-HSA-422475), hemostasis (R-HSA-109582), and cell surface interactions at the vascular wall (R-HSA-202733) (Fig. 4C). Although macrophages must be able to pass blood vessel walls and have been implicated in clotting, the last two pathways were mostly included in the others (Fig. S7). Moreover, due to the *in vitro* setting of the dataset, we disregarded hemostasis and interactions at the vascular wall and focused on the first six pathways. Figure 4D shows the pathway-annotated network.

We identified a platform of LRIs involved collagens (COL4A1, COL5A2, COL6A2, COL6A3) and integrins (ITGB1, ITGB5, ITGAV) as well as CD47 linked to adaptive immunity and neutrophil degranulation, two functions that are increased in differentiated macrophages. Moreover, incubation of U-937 cells with PMA increases adherence and the expression of integrin β2 (ITGB2) [48]. Indeed, the expression of ITGB2 was increased as well as that of ITGB1, ITGB5, and ITGAV. ECM remodeling increases during monocyte-to-macrophage differentiation [49]. In line, the macrophage network contained many collagen and metalloproteinase molecules targeted by integrin-triggered pathways. ECM remodeling also relates to another identified LRI (*i.e.*, VIM-CD44 interaction) that is common and highly conserved in circulating immune cells [50]. RTK signaling involved ITGB1 and ITGAV as receptors, although they are no RTKs. Therefore, this pathway annotation is more relevant for the set of downstream molecules that might be activated by PMA through mitogen-activated protein kinase networks in macrophages [51–53]. RTK signaling was strongly linked to adaptive immunity in the macrophage network.

The muscle contraction and the neural system pathways were also activated, both mainly triggered by the calmodulin 3 (CALM3)-potassium voltage-gated channel subfamily Q member 1 (KCNQ1) interaction, and the various ligands of ITGB5. The muscle contraction subnetwork combined myosin light and heavy chain proteins (MYH6, MYH11, MYL1, MYL6, MYL7, MYL12A) and proteins involved in ion exchanges, such as calcium voltage-gated channel complexes (*e.g.*, CACNA2D2, CACNA1F, CACNG6), sodium voltage-gated channels (*e.g.*, SCNA4, SCN11A), calcium/sodium exchangers (*e.g.*, SLC8A2, SLS8A3), and cell internal ion transporters (*e.g.*, ATP1A4, ATP2A2, ATP2B3, ATP2B4). The neural system-annotated proteins were involved in potassium, calcium and sodium transport. This network also contained the enzyme glutamate-ammonia ligase (GLUL) that catalyzes the synthesis of glutamine from glutamate. SLC1A1, a high-affinity glutamate transporter family member was upstream of GLUL in the network. This last observation might be related to the ability of macrophages to produce glutamine for other cells [54]. A number of additional genes were annotated in the axon guidance pathway, which mostly related to focal adhesion and further motility (MYH9, MYH10, MYH14, MYO10, ARPC1B, ARPC2, ARPC3, ARPC4). Altogether, these pathways reflected increased motility and might be associated with the management of ion fluxes.

We also identified two LRIs that were not included in any of the six main pathways and that involved G protein subunit alpha i2 (GNAI2) and were related to G protein-coupled receptor (GPCR) signaling: GNAI2-complement C5a receptor 1 (C5AR1) and GNAI2-dopamin receptor D2 (DRD2). GPCR signaling in macrophages has many roles, including sensing inflammatory molecules (*e.g.*, chemokines) and also the C5 cleavage product C5a through C5AR1 [55]. The GNAI2-DRD2 interaction regulates DRD2 activity by limiting its access to the membrane [56] and DRD2 controls inflammation in macrophages [57,58].

### Complementarity with transcriptomic data

The authors of the above scProt-MS dataset also provided SCT data obtained from untreated and PMA-treated U-937 cells. We used these SCT data to perform LRI analysis and to compare the obtained proteomic- and transcriptomic-based results. The SCT data covered 20,274 cells and 32,738 genomic features. After filtering low-quality transcriptomes and undetected features (Materials and Methods), we obtained a transcript expression matrix for 4,296 genes in 7,177 cells (Fig. S2A), including 113 ligands and 91 receptors. Cells were sorted according to the average *VIM*, *ITGB2*, elastase neutrophil expressed (*ELANE*), and nucleolin (*NCL*) expression to define the top 20% cells (macrophages), and bottom 20% cells (monocytes) (Fig. S2B-C). *ELANE* and *NCL* were considered macrophage markers in the SCT data by the dataset authors [18].

We identified 29 ligands and 17 receptors the expression of which significantly increased in macrophages and that were involved in 42 significant LRIs (Supplementary Data). Figure 5A shows the parsimonious network linking the identified transcriptomic-based LRIs with their downstream targets (in Fig. S8 with gene names). Several biological pathways associated with the receptors were common with those identified in the proteomic analysis, such as ECM remodeling and adaptive immune system. Conversely, motility-related pathways were much reduced. Moreover, signaling was triggered by different sets of receptors, while for downstream signaling we found shared and specific molecules. Signaling by RTK was much reduced, but newly identified pathways such as interleukin signaling (R-HSA-449147) and GPCR signaling (G alpha (i) signalling events, R-HSA-418594). We also identified the hemostasis and cell surface interactions at the vascular wall pathways, but with very few or no specific genes (Fig. S9). We decided to ignore them in our analysis. Lastly, a number of network nodes fall in no aforementioned pathway, the main part of which were annotated as signaling by nuclear receptors (R-HSA-9006931) and oncogenic MAPK signaling (R-HSA-6802957).

**Figure 5.**
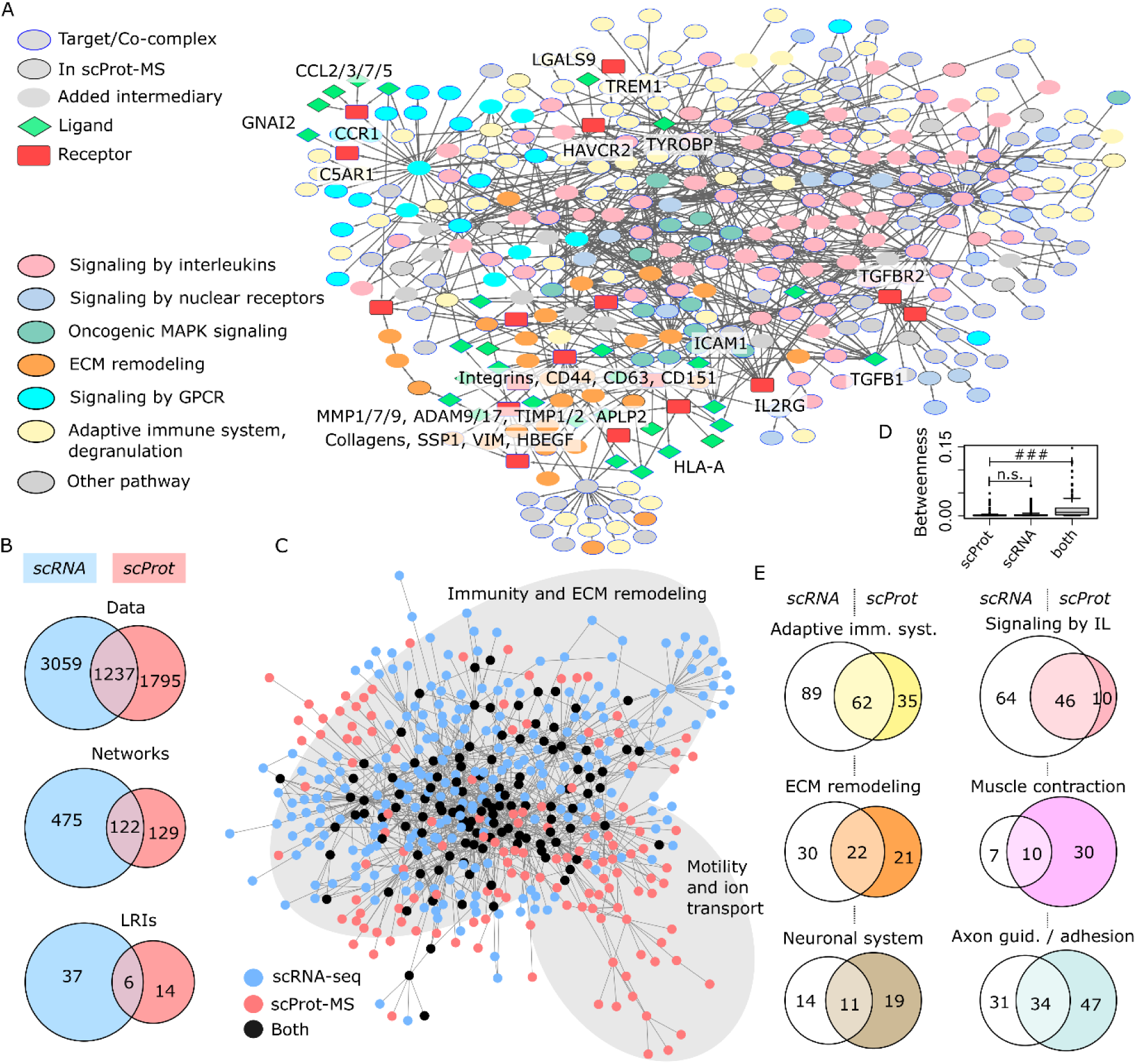
Comparison with scRNA-seq data. (**A**) The macrophage autocrine LRI network revealed using scRNA-seq data. (**B**) Distinct and overlapping features of scRNA-seq and scProt-MS data, the corresponding LRI networks, and LRIs. (**C**) Merge of the scRNA-seq and scProt-MS LRI networks. Shared and distinct modules are highlighted. (**D**) Comparison of betweenness centrality of nodes detected in distinct modalities and shared nodes (### stands for P < 2.2E-16, Wilcoxon test). (**E**) Number of nodes contributed by each modality in selected pathways.

Most of the obtained network relates immunity, ECM remodeling, and various signaling pathways. GPCR signaling, which was much weaker in the proteomic-based analysis and essentially limited to the GNAI2-C5AR1 LRI, was predominant here, in line with its known importance in macrophages [55,59]. Several LRIs were related to inflammation, such as interleukin signaling that was triggered by the intercellular adhesion molecule 1 (ICAM1)-interleukin 2 receptor γ (IL2RG) interaction [60], or LRIs involving several integrins (ITGB1, ITGB2, ITGAX). Additional LRIs linked CC motif chemokine receptor 1 (CCR1) to chemokine ligand 2, 3, 5, and 7 (CCL2/3/5/7). Moreover, the interaction between transmembrane immune signaling adaptor (TYROBP) and triggering receptor expressed on myeloid cells 1 (TREM1) facilitates TREM1 multimerization and activation as a mediator of inflammation [61]. We also found the galectin 9 (LGALS9)-hepatitis a virus cellular receptor (HAVCR2, TIM-3) interaction, which is an immune checkpoint, thereby indicating the ability to counterweigh spontaneous, autocrine inflammatory signals by this autocrine checkpoint in the absence of pathogens.

The interaction between major histocompatibility complex, class I (HLA-A) and amyloid beta precursor like protein 2 (APLP2) is related to the MHC Class I molecule cell surface expression regulation [62]. Its autocrine nature suggests that macrophages can keep the expression of MHC class I molecules at their surface at a basal level until the same mechanism in an inflammatory context will adjust their expression. The interaction of tumor growth factor β1 (TGFB1) with TGF-β receptor 2 (TGFBR2) induces signals that can trigger various cell responses, such as proliferation, differentiation, motility, and apoptosis. The expression levels of *TGFB1* and *TGFBR2* are increased in several PMA-treated monocyte models [63], and autocrine TGF-β is a survival factor for CD14+ monocytes that might polarize macrophage differentiation [64].

Although the transcriptomics data did not directly point towards motility-related pathways, two transmembrane 4 superfamily (tetraspanin family) members, CD63 [65,66] and CD151 [67,68], are associated with motility and metastasis in tumors. They interacted with metallopeptidase inhibitor 1 (TIMP1) and matrix metalloproteinase-7 (MMP7), respectively.

These results suggest complementarity between modalities, although the scRNA-seq dataset had higher coverage depth. Indeed, the two modalities covered rathe different molecules (Fig. 5B top, nonsignificant intersection, P = 0.34, hypergeometric test), which led to two significantly different sets of LRIs (Fig. 5B bottom, P = 0) that nonetheless functionally converged to networks with a significant overlap (Fig. 5B middle, P = 1.7E-79). Figure 5C shows the shared and distinct molecules in the proteomic- and transcriptomic-based networks. Molecules identified by the two modalities were more central (Fig. 5D). Inspection of the merged network in Figure 5C revealed a large shared part with immunity and ECM remodeling accompanied with different types of signaling, and a proteomic-specific part enriched in pathways related to motility and ion transport. Comparing scRNA-seq- and scProt-MS-identified molecules in various pathways (Fig. 5E), we observed contributions commensurate with dataset sizes in immunity, an enrichment of interleukin signaling in scRNA-seq, and an enrichment in ECM remodeling and motility in scProt-MS.

### Application to bulk analysis of FACS-sorted cell populations

Choi, *et al*. [22] studied a mouse non-small cell lung cancer (NSCLC) model (*Kras*^G12D/+^, *p53*^-/-^) and its microenvironment. They FACS sorted epithelial cells, neutrophils, monocytes, and macrophages from tumors and WT animals (2 or 3 replicates for each population/condition) and submitted to RNA sequencing. For every cell population, the comparison of the tumor and matching WT samples identified genes whose transcripts were deregulated in cancer. Thanks to our new framework, we could define our own function to perform the same comparisons using edgeR that is perfectly appropriate in this classical RNA-sequencing setting. Choi, *et al*. were interested in ligands secreted by the three immune populations and their signaling impact on cancer epithelial cells. They used an algorithm (CCCExplorer) that is precursor of most LRI inference methods using pathways and intracellular molecular interactions.

Globally, we found the same interactions (Supplemental Material) as in the original work. They put a particular emphasis on interleukin inflammatory signals and the macrophage-epithelial cell IL6-IL6R interaction, which they showed to activate STAT3 *in vitro*. Starting from a more global perspective, we found a lot of LRIs between cell populations that were deregulated in cancer (Fig. 6A). Interleukin signaling was ubiquitously augmented reflecting an inflammatory microenvironment (Fig. 6B).

**Figure 6.**
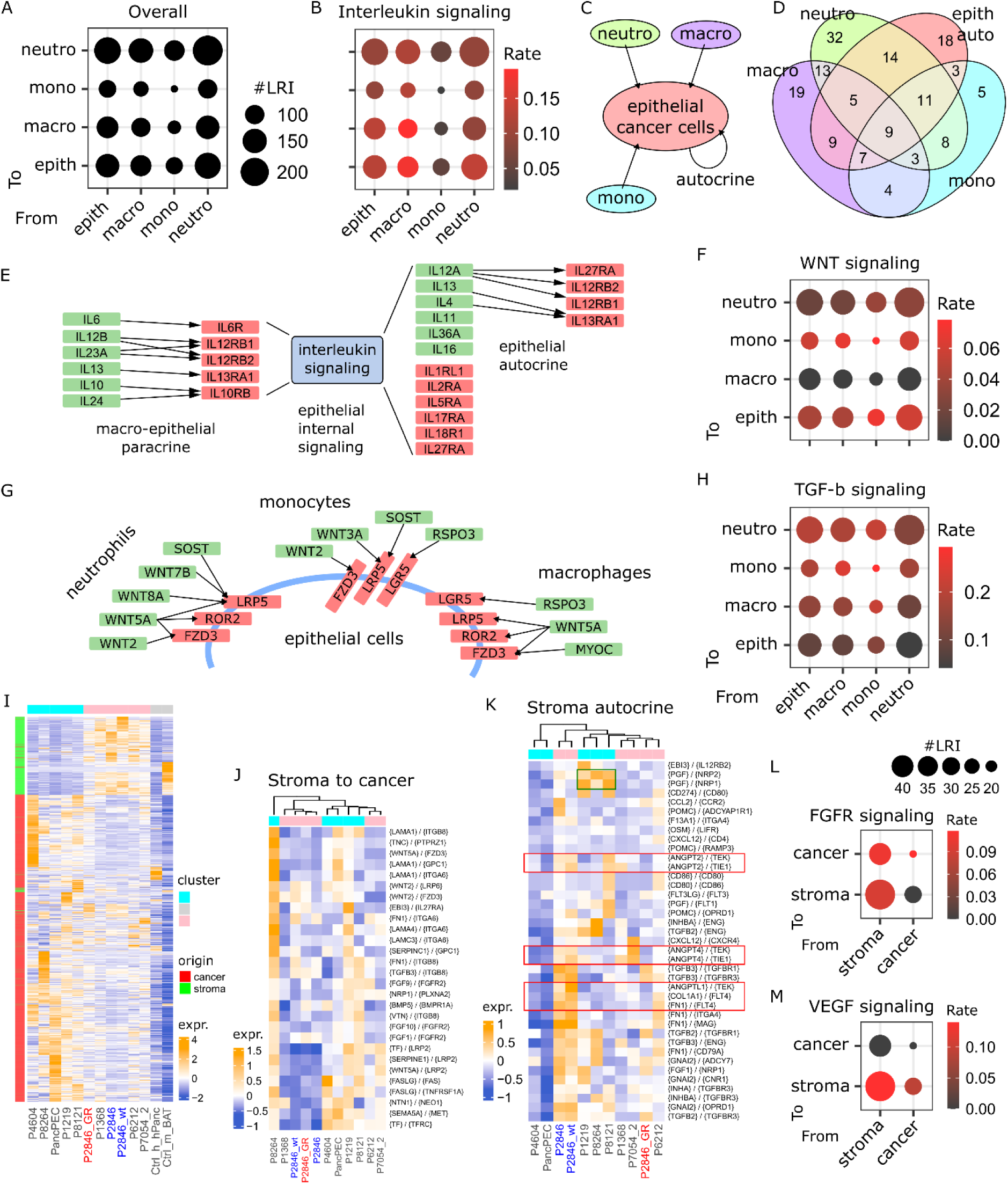
Additional applications. (**A**) Overview of LRI numbers between selected NSCLC tumor populations. (**B**) Proportion of interleukin signaling. (**C**) Total LRIs at epithelial cancer cells. (**D**) Diversity of ligands involved in LRIs toward epithelial cancer cells. (**E**) Macrophage-cancer cell and cancer cell autocrine interleukin LRIs. (**F**) Proportion of Wnt signaling. (**G**) Paracrine Wnt signaling towards epithelial cancer cells. (**H**) Proportion of TGF-β signaling. (**I**) Overview of PDX transcriptomes. The original P2846 model (wt, blue) was made germcitabine resistant (GR, red), and was also available untreated after the same number of passes as the GR model (P2648, blue). (**J**) Scoring of stroma to cancer cell paracrine LRIs across all PDX. (**K**) Scoring of stromal autocrine LRIs. Green square = angiogenesis-related LRIs. Red rectangles = angiogenesis- and lymphoangiogenesis-related LRIs that are reduced in P2846_GR compared to P2846 and P2846_wt. (**L**) FGFR signaling. (**M**) VEGF signaling.

Focusing on signals towards cancer cells, including cancer autocrine LRIs (Fig. 6C), we found that cancer epithelial cells were subjected to a multitude of shared and specific deregulated ligands (Fig. 6D). We reduced the analysis to interleukin signaling from macrophages to cancer cells, including cancer autocrine interleukin signals (Fig. 6E). We found that cancer cell intracellular interleukin signaling triggered by macrophage ligands lead to the expression of variety of deregulated interleukin ligands (or their subunits) that were different from those secreted by macrophages: IL4, IL12A, IL11, IL16, and IL36A. Part of these ligands were involved in autocrine signals (IL4, IL12A, IL13). The other ligands could interact with the microenvironment as suggested by Figure 6B. Interleukin receptors that did not participate to interactions with macrophages were also upregulated: IL1RL1, IL2RA, IL5RA, IL17RA, IL18R1, and IL27RA. Compared to the original publication, many more interleukins were found here though this could be related to thresholds in differential expression analysis.

Switching to Wnt signaling that is known to play an important role in the lung [69] and lung cancer [70], we observed a different pattern (Fig. 6F). Ligands still originated from the four cell populations, but their receptors were mostly expressed at cancer epithelial cells and cancer-associated monocytes. Interestingly, concentrating on cancer epithelial cells, we found that the three immune populations contributed slightly different ligand sets targeting both the canonical and noncanonical Wnt pathways (Fig. 6G). We also note that regulators Wnt signaling, both positive (RSPO3, LGR5, LPR5) and negative (SOST, ROR2), as well as associated genes (MYOC) were identified. Actual regulation at the protein level might differ.

One last example of signal that adopted yet another configuration was TGF-β (Fig. 6H). It was less pronounced towards cancer cells and stronger towards the stroma. This is coherent with a role that is commonly associated to the extracellular matrix, fibroblast reprogramming, and immunosuppression.

### Application to PDX bulk RNA-sequencing data

In PDX, the stromal compartment consists of murine cells, whereas the cancer compartment is human. Bulk transcriptomic profiling thus allows separation of the two components by aligning reads against the respective genomes [23]. Application of SingleCellSignalR enables mapping of LRIs between these tumor compartments.

To illustrate this methodology, we analyzed 11 PDAC PDX [40,41], complemented with healthy human pancreas and mouse brown adipose tissue as control samples. Alignment against the two genomes identified 10,519 cancer genes and 2,873 stromal genes (Fig. S10), whose expression profiles are featured in Figure 6I. Two PDX clusters emerged, distinguished by the abundance of stromal transcripts. As PDX were grown in nude mice, the stroma mainly consisted of cancer-associated fibroblasts (CAFs) and neovessels, with minimal immune infiltration. The P2846 PDX was available in three versions: unmodified (P2846_wt, 3 passages), gemcitabine-resistant (P2846_GR, 7 passages), and a passage-matched untreated control (P2846) [41]. As shown in Figure 6I, the P2846_GR PDX displayed some differences compared to P2846 and P2846_wt.

By exploiting SingleCellSignalR v2 framework as for FACS-separated cells above, we implemented a tailored function for differential gene expression analysis. Human and mouse transcripts were combined into a single expression matrix, with differential analyses performed separately by comparing human genes in PDX to the control human cell line, and murine genes to the murine control tissue. Figure 6J shows LRI activity scores for stroma-to-cancer LRIs. These scores were computed using BulkSignalR [30] and represent a weighted average of the ligand, receptor and target genes normalized expression. This analysis revealed that LRI activity did not simply mirror the rich/poor stroma clusters, as both groups contained PDX with high or low scores. No marked differences were observed among the three P2846 models. In contrast, stromal autocrine interactions related to angio- and lymphoangiogenesis displayed markedly reduced scores in P2846_GR PDX (Fig. 6K). Given that gemcitabine has been reported to modulate angiogenesis in PDAC [71,72], this finding may suggest resistance through altered neovascularization in the GR PDX. Moreover, three PDX (P1219, P8264, and P8121) exhibited distinct angiogenesis-related LRIs involving placental growth factor (PGF)–neuropilin 1 (NRP1) and PGF–NRP2, consistent with recent evidence that PGF/VEGF blockade is relevant for PDAC murine models [73]. Activity scores obtained for cancer-to-stroma and cancer autocrine LRIs are reported in Figure S11.

As expected in the context of nude mice lacking a functional immune system, stromal autocrine LRIs were largely associated with angiogenesis, cell contacts and extracellular matrix organization. Moreover, we observed that fibroblast growth factors (FGFs) were secreted predominantly by stromal cells and mainly targeted cancer cells, although some stromal autocrine signaling was also detected (Fig. 6L). VEGF signaling displayed a distinct architecture, with the stroma acting as the exclusive target. In this case, secretion was largely confined to stromal cells, with minimal contribution from cancer cells (Fig. 6M).

## DISCUSSION AND CONCLUSIONS

The inference of LRIs from single-cell expression data is a well-recognized yet multifaceted problem. Outcomes depend strongly on the experimental design, the biological question posed, and the data modality considered, whether transcriptomic, proteomic, ATAC-seq, spatial, or even multimodal. This diversity, combined with the creativity of researchers, has led to the development of nearly 100 dedicated tools [2]. Here, we contributed an expanded version of our tool SingleCellSignalR [9] that addresses the most common use case that is SCT and, since recently, scProt-MS.

Version 2 of our tool introduces a new statistical model for scoring LRIs based on the differential expression of ligands, receptors, and their target genes downstream Reactome pathways and GO BP terms. This comparative approach, termed SCSRNet, is designed to identify LRIs with enhanced activity in specific cell populations. However, in settings where cell populations display globally similar profiles, differential approaches may obscure relevant interactions. To account for such cases, we retained the original expression-based scoring scheme (LR-score), here named SCSRNoNet. Its utility was highlighted on a dataset of closely related populations, where expression-based tools (SCSRNoNet, CytoTalk [34], and cellChat [31]) successfully recovered the expected LRI, while differential schemes (SCSRNet, scMLnet [33], and scSeqComm [32]) failed.

To evaluate performance in a more conventional setting, we analyzed two comparable respiratory airway cell atlases from HLCA [39]. This choice provided both population diversity and the opportunity to assess reproducibility across comparable datasets generated in distinct laboratories. As expected, SCSRNoNet detected more LRIs than the stricter SCSRNet, which additionally requires evidence of downstream target regulation; notably, nearly all SCSRNet inferences were included within those of SCSRNoNet. Across both atlases, SCSRNoNet was the most sensitive method, whereas CytoTalk was the least. SCSRNet was more selective than scMLnet, scSeqComm, and CellChat, with reproducibility across atlases comparable to these methods. Importantly, SCSRNoNet achieved the highest reproducibility in this comparison.

In the absence of a definitive ground truth, and given the large number of available tools, we did not attempt a comprehensive benchmark. Previous studies [3,4] have shown that tool outputs often diverge considerably, suggesting that all current methods likely produce false positives (FPs) while also missing genuine interactions. Based on our tests, we simply concluded that our framework provides a versatile solution: SCSRNoNet as a highly sensitive, universal option (albeit more FP-prone), and SCSRNet as a stricter, differential approach that benefits from receptor downstream modeling and is likely less FP-prone. In practice, SCSRNet offers numerous adjustable parameters (not detailed here) controlling the extent of evidence required for receptor signaling, as well as the depth of pathway exploration, enabling users to balance sensitivity and specificity. We consider such flexibility, coupled with visualization functions, essential for exploring LRIs in diverse experimental settings.

Our main example application concerned scProt-MS. Most scProt-MS datasets generated until recently achieved a depth of ∼1,100-1,300 proteins [14,44,74–76]. At this coverage, ligand and receptor expression data were sufficient to identify cell populations, but not for mapping LRIs. We thus selected a deeper dataset reporting 3,032 proteins from U937 monocytes, untreated or incubated with PMA to induce their differentiation into macrophages [18]. Matching SCT data were also available, allowing direct comparison of the application of SCSRNet to both modalities. Since untreated monocytes and differentiated macrophages were cultured separately, paracrine LRIs were irrelevant. We inferred autocrine LRIs enriched in macrophages. From scProt-MS, we found LRIs associated with functions consistent with the acquired macrophage phenotype, including immune system, motility, RTK signaling, and ECM remodeling. From SCT data, we recovered LRIs and pathways that significantly overlapped those found using scProt-MS data (interleukin signaling, immune system, ECM remodeling, GPCR signaling). Global statistics further supported this observation: despite relying on significantly disjoint sets of genes versus proteins, the two modalities yielded significantly overlapping molecular networks. Notably, scProt-MS data uniquely highlighted a subnetwork related to motility and potentially associated ion transport, which was much less prevalent in SCT, underscoring the complementarity of the two modalities.

Leveraging the flexibility of SingleCellSignalR version 2, we further demonstrated its ability to handle common yet atypical data types in the LRI field. In a first example, we analyzed bulk transcriptomes of targeted, FACS-sorted cell populations. We used a NSCLC dataset comprising epithelial cancer cells and multiple immune cell types [22]. Compared to the original analysis, SCSRNet provided deeper insight into intercellular communication. In a second example, we showed how preclinical PDX models can be explored with SCSRNet to learn about LRIs both across and within their cancer and stromal compartments. To our knowledge, no other LRI inference tool currently supports this type of analysis, underscoring the versatility of our framework.

Altogether, the expanded version of SingleCellSignalR, built upon the models and functionalities of its sister package BulkSignalR [30], offers multiple options to explore LRIs across diverse experimental settings. The parallel analyses of paired scProt-MS and SCT data highlighted the value of a versatile and integrated framework, while the application to FACS-sorted bulk transcriptomes and PDX models further underscored its ability to adapt seamlessly to atypical data types.

## ACKNOWLEDGMENTS

JC was supported by grants INCa PRT-K 2020-038 and ANR 21-CE13-0011-03.

## DATA AND CODE AVAILABILITY

All the data were publicly available (Materials and Methods) except PDX data that were submitted to GEO. SingleCellSignalR is available from Bioconductor. Version 2 is included in the devel version at the time of writing. It is also available from GitHub directly at https://github.com/jcolinge/SingleCellSignalR. The R code used to process the three detailed examples is available from SingleCellSignalR companion GitHub repository https://github.com/jcolinge/SingleCellSignalR_companion.

## SUPPLEMENTARY DATA

Upon publication.

## REFERENCES

1. Wang X, Almet AA, Nie Q. The promising application of cell-cell interaction analysis in cancer from single-cell and spatial transcriptomics. Semin. Cancer Biol. 2023; 95:42–51

2. Armingol E, Baghdassarian HM, Lewis NE. The diversification of methods for studying cell-cell interactions and communication. Nat. Rev. Genet. 2024; 25:381–400

3. Dimitrov D, Türei D, Garrido-Rodriguez M, et al. Comparison of methods and resources for cell-cell communication inference from single-cell RNA-Seq data. Nat. Commun. 2022; 13:3224

4. Liu Z, Sun D, Wang C. Evaluation of cell-cell interaction methods by integrating single-cell RNA sequencing data with spatial information. Genome Biol. 2022; 23:218

5. Shaw R, Tian X, Xu J. Single-Cell Transcriptome Analysis in Plants: Advances and Challenges. Mol. Plant 2021; 14:115–126

6. Stuart T, Satija R. Integrative single-cell analysis. Nat. Rev. Genet. 2019; 20:257–272

7. Cha J, Lee I. Single-cell network biology for resolving cellular heterogeneity in human diseases. Exp. Mol. Med. 2020; 52:1798–1808

8. Schwartzman O, Tanay A. Single-cell epigenomics: techniques and emerging applications. Nat. Rev. Genet. 2015; 16:716–726

9. Cabello-Aguilar S, Alame M, Kon-Sun-Tack F, et al. SingleCellSignalR: inference of intercellular networks from single-cell transcriptomics. Nucleic Acids Res. 2020;

10. Sahni N, Yi S, Taipale M, et al. Widespread Macromolecular Interaction Perturbations in Human Genetic Disorders. Cell 2015; 161:647–660

11. Luck K, Kim D-K, Lambourne L, et al. A reference map of the human binary protein interactome. Nature 2020; 580:402–408

12. Khan Z, Ford MJ, Cusanovich DA, et al. Primate Transcript and Protein Expression Levels Evolve Under Compensatory Selection Pressures. Science 2013; 342:1100–1104

13. Buchan DW, Rison SC, Bray JE, et al. Classical Hodgkin’s lymphoma in adults: guidelines of the Italian Society of Hematology, the Italian Society of Experimental Hematology, and the Italian Group for Bone Marrow Transplantation on initial work-up, management, and follow-up. Proc. Natl. Acad. Sci. U. S. A. 2013; 31:7158–63

14. Budnik B, Levy E, Harmange G, et al. SCoPE-MS: mass spectrometry of single mammalian cells quantifies proteome heterogeneity during cell differentiation. Genome Biol. 2018; 19:161

15. Petrosius V, Schoof EM. Recent advances in the field of single-cell proteomics. Transl. Oncol. 2023; 27:101556

16. Ahmad R, Budnik B. A review of the current state of single-cell proteomics and future perspective. Anal. Bioanal. Chem. 2023; 415:6889–6899

17. Furtwängler B, Üresin N, Richter S, et al. Mapping early human blood cell differentiation using single-cell proteomics and transcriptomics. Science 2025; eadr8785

18. Specht H, Emmott E, Petelski AA, et al. Single-cell proteomic and transcriptomic analysis of macrophage heterogeneity using SCoPE2. Genome Biol. 2021; 22:50

19. Khan S, Elcheikhali M, Leduc A, et al. Inferring post-transcriptional regulation within and across cell types in human testis. 2024; 2024.10.08.617313

20. Woo J, Williams SM, Markillie LM, et al. High-throughput and high-efficiency sample preparation for single-cell proteomics using a nested nanowell chip. Nat. Commun. 2021; 12:6246

21. Marx V. A dream of single-cell proteomics. Nat. Methods 2019; 16:809–812

22. Choi H, Sheng J, Gao D, et al. Transcriptome analysis of individual stromal cell populations identifies stroma-tumor crosstalk in mouse lung cancer model. Cell Rep. 2015; 10:1187–1201

23. Rossello FJ, Tothill RW, Britt K, et al. Next-generation sequence analysis of cancer xenograft models. PloS One 2013; 8:e74432

24. Kanehisa M, Goto S. KEGG: kyoto encyclopedia of genes and genomes. Nucleic Acids Res. 2000; 28:27–30

25. Gillespie M, Jassal B, Stephan R, et al. The reactome pathway knowledgebase 2022. Nucleic Acids Res. 2022; 50:D687–D692

26. Rodchenkov I, Babur O, Luna A, et al. Pathway Commons 2019 Update: integration, analysis and exploration of pathway data. Nucleic Acids Res. 2020; 48:D489–D497

27. The Gene Ontology Consortium, Aleksander SA, Balhoff J, et al. The Gene Ontology knowledgebase in 2023. Genetics 2023; 224:iyad031

28. Müller-Dott S, Tsirvouli E, Vazquez M, et al. Expanding the coverage of regulons from high-confidence prior knowledge for accurate estimation of transcription factor activities. Nucleic Acids Res. 2023; 51:10934–10949

29. Liberzon A, Subramanian A, Pinchback R, et al. Molecular signatures database (MSigDB) 3.0. Bioinformatics 2011; 27:1739–1740

30. Villemin J-P, Bassaganyas L, Pourquier D, et al. Inferring ligand-receptor cellular networks from bulk and spatial transcriptomic datasets with BulkSignalR. Nucleic Acids Res. 2023; gkad352

31. Jin S, Guerrero-Juarez CF, Zhang L, et al. Inference and analysis of cell-cell communication using CellChat. Nat. Commun. 2021; 12:1088

32. Baruzzo G, Cesaro G, Di Camillo B. Identify, quantify and characterize cellular communication from single-cell RNA sequencing data with scSeqComm. Bioinforma. Oxf. Engl. 2022; 38:1920–1929

33. Cheng J, Zhang J, Wu Z, et al. Inferring microenvironmental regulation of gene expression from single-cell RNA sequencing data using scMLnet with an application to COVID-19. Brief. Bioinform. 2021; 22:988–1005

34. Hu Y, Peng T, Gao L, et al. CytoTalk: De novo construction of signal transduction networks using single-cell transcriptomic data. Sci. Adv. 2021; 7:eabf1356

35. Wilk AJ, Shalek AK, Holmes S, et al. Comparative analysis of cell–cell communication at single-cell resolution. Nat. Biotechnol. 2024; 42:470–483

36. Clough E, Barrett T, Wilhite SE, et al. NCBI GEO: archive for gene expression and epigenomics data sets: 23-year update. Nucleic Acids Res. 2023; 52:D138–D144

37. Hao Y, Stuart T, Kowalski MH, et al. Dictionary learning for integrative, multimodal and scalable single-cell analysis. Nat. Biotechnol. 2024; 42:293–304

38. Aran D, Looney AP, Liu L, et al. Reference-based analysis of lung single-cell sequencing reveals a transitional profibrotic macrophage. Nat. Immunol. 2019; 20:163–172

39. Sikkema L, Ramírez-Suástegui C, Strobl DC, et al. An integrated cell atlas of the lung in health and disease. Nat. Med. 2023; 29:1563–1577

40. Bruciamacchie M, Garambois V, Vie N, et al. ATR inhibition potentiates FOLFIRINOX cytotoxic effect in models of pancreatic ductal adenocarcinoma by remodelling the tumour microenvironment. Br. J. Cancer 2024;

41. Rabia E, Garambois V, Hubert J, et al. Anti-tumoral activity of the Pan-HER (Sym013) antibody mixture in gemcitabine-resistant pancreatic cancer models. mAbs 2021; 13:1914883

42. Giguelay A, Turtoi E, Khelaf L, et al. The landscape of cancer-associated fibroblasts in colorectal cancer liver metastases. Theranostics 2022; 12:7624–7639

43. Kieffer Y, Hocine HR, Gentric G, et al. Single-Cell Analysis Reveals Fibroblast Clusters Linked to Immunotherapy Resistance in Cancer. Cancer Discov. 2020; 10:1330–1351

44. Schoof EM, Furtwängler B, Üresin N, et al. Quantitative single-cell proteomics as a tool to characterize cellular hierarchies. Nat. Commun. 2021; 12:3341

45. Collins BC, Hunter CL, Liu Y, et al. Multi-laboratory assessment of reproducibility, qualitative and quantitative performance of SWATH-mass spectrometry. Nat. Commun. 2017; 8:291

46. Gebreyesus ST, Siyal AA, Kitata RB, et al. Streamlined single-cell proteomics by an integrated microfluidic chip and data-independent acquisition mass spectrometry. Nat. Commun. 2022; 13:37

47. Leduc A, Huffman RG, Cantlon J, et al. Exploring functional protein covariation across single cells using nPOP. Genome Biol. 2022; 23:261

48. García A, Serrano A, Abril E, et al. Differential effect on U937 cell differentiation by targeting transcriptional factors implicated in tissue-or stage-specific induced integrin expression. Exp. Hematol. 1999; 27:353–364

49. Chang MY, Chan CK, Braun KR, et al. Monocyte-to-macrophage differentiation: synthesis and secretion of a complex extracellular matrix. J. Biol. Chem. 2012; 287:14122–14135

50. Li Z, Sun C, Wang F, et al. Molecular mechanisms governing circulating immune cell heterogeneity across different species revealed by single-cell sequencing. Clin. Transl. Med. 2022; 12:e689

51. Richter E, Ventz K, Harms M, et al. Induction of Macrophage Function in Human THP-1 Cells Is Associated with Rewiring of MAPK Signaling and Activation of MAP3K7 (TAK1) Protein Kinase. Front. Cell Dev. Biol. 2016; 4:21

52. Traore K, Trush MA, George M, et al. Signal transduction of phorbol 12-myristate 13-acetate (PMA)-induced growth inhibition of human monocytic leukemia THP-1 cells is reactive oxygen dependent. Leuk. Res. 2005; 29:863–879

53. Qiu ZH, Leslie CC. Protein kinase C-dependent and -independent pathways of mitogen-activated protein kinase activation in macrophages by stimuli that activate phospholipase A2. J. Biol. Chem. 1994; 269:19480–19487

54. Shang M, Cappellesso F, Amorim R, et al. Macrophage-derived glutamine boosts satellite cells and muscle regeneration. Nature 2020; 587:626–631

55. Lavallée-Adam M, Cloutier P, Coulombe B, et al. Anti-HER3 domain 1 and 3 antibodies reduce tumor growth by hindering HER2/HER3 dimerization and AKT-induced MDM2, XIAP, and FoxO1 phosphorylation. Int. J. Radiat. Oncol. Biol. Phys. 2013; 9 Suppl 9:e1000217

56. López-Aranda MF, Acevedo MJ, Gutierrez A, et al. Role of a Galphai2 protein splice variant in the formation of an intracellular dopamine D2 receptor pool. J. Cell Sci. 2007; 120:2171–2178

57. Han X, Ni J, Wu Z, et al. Myeloid-specific dopamine D2 receptor signalling controls inflammation in acute pancreatitis via inhibiting M1 macrophage. Br. J. Pharmacol. 2020; 177:2991–3008

58. Nolan RA, Muir R, Runner K, et al. Role of Macrophage Dopamine Receptors in Mediating Cytokine Production: Implications for Neuroinflammation in the Context of HIV-Associated Neurocognitive Disorders. J. Neuroimmune Pharmacol. 2019; 14:134–156

59. Wang X, Iyer A, Lyons AB, et al. Emerging Roles for G-protein Coupled Receptors in Development and Activation of Macrophages. Front. Immunol. 2019; 10:2031

60. Burton J, Goldman CK, Rao P, et al. Association of intercellular adhesion molecule 1 with the multichain high-affinity interleukin 2 receptor. Proc. Natl. Acad. Sci. U. S. A. 1990; 87:7329–7333

61. Carrasco K, Boufenzer A, Jolly L, et al. TREM-1 multimerization is essential for its activation on monocytes and neutrophils. Cell. Mol. Immunol. 2019; 16:460–472

62. Tuli A, Sharma M, Naslavsky N, et al. Specificity of amyloid precursor-like protein 2 interactions with MHC class I molecules. Immunogenetics 2008; 60:303–313

63. Riddy DM, Goy E, Delerive P, et al. Comparative genotypic and phenotypic analysis of human peripheral blood monocytes and surrogate monocyte-like cell lines commonly used in metabolic disease research. PloS One 2018; 13:e0197177

64. Gonzalez-Junca A, Driscoll KE, Pellicciotta I, et al. Autocrine TGFβ Is a Survival Factor for Monocytes and Drives Immunosuppressive Lineage Commitment. Cancer Immunol. Res. 2019; 7:306–320

65. Radford KJ, Thorne RF, Hersey P. Regulation of tumor cell motility and migration by CD63 in a human melanoma cell line. J. Immunol. Baltim. Md 1950 1997; 158:3353–3358

66. Lupia A, Peppicelli S, Witort E, et al. CD63 tetraspanin is a negative driver of epithelial-to-mesenchymal transition in human melanoma cells. J. Invest. Dermatol. 2014; 134:2947–2956

67. Palmer TD, Martínez CH, Vasquez C, et al. Integrin-Free Tetraspanin CD151 Can Inhibit Tumor Cell Motility upon Clustering and Is a Clinical Indicator of Prostate Cancer Progression. Cancer Res. 2014; 74:173–187

68. Hong I-K, Jin Y-J, Byun H-J, et al. Homophilic interactions of Tetraspanin CD151 up-regulate motility and matrix metalloproteinase-9 expression of human melanoma cells through adhesion-dependent c-Jun activation signaling pathways. J. Biol. Chem. 2006; 281:24279–24292

69. Aros CJ, Pantoja CJ, Gomperts BN. Wnt signaling in lung development, regeneration, and disease progression. Commun. Biol. 2021; 4:601

70. Zhu W, Wang H, Zhu D. Wnt/β-catenin signaling pathway in lung cancer. Med. Drug Discov. 2022; 13:100113

71. Yang Y, Tian W, Yang L, et al. Gemcitabine potentiates anti-tumor effect of resveratrol on pancreatic cancer via down-regulation of VEGF-B. J. Cancer Res. Clin. Oncol. 2021; 147:93–103

72. Khan MA, Srivastava SK, Bhardwaj A, et al. Gemcitabine triggers angiogenesis-promoting molecular signals in pancreatic cancer cells: Therapeutic implications. Oncotarget 2015; 6:39140– 39150

73. Kim DK, Jeong J, Lee DS, et al. PD-L1-directed PlGF/VEGF blockade synergizes with chemotherapy by targeting CD141+ cancer-associated fibroblasts in pancreatic cancer. Nat. Commun. 2022; 13:6292

74. Fulcher JM, Markillie LM, Mitchell HD, et al. Parallel measurement of transcriptomes and proteomes from same single cells using nanodroplet splitting. 2022; 2022.05.17.492137

75. Petrosius V, Aragon-Fernandez P, Üresin N, et al. Exploration of cell state heterogeneity using single-cell proteomics through sensitivity-tailored data-independent acquisition. Nat. Commun. 2023; 14:5910

76. Derks J, Leduc A, Wallmann G, et al. Increasing the throughput of sensitive proteomics by plexDIA. Nat. Biotechnol. 2023; 41:50–59

