## Supplemental information for "Addressing multiple facets of ligand-receptor network inference including single-cell proteomics"

### SUPPLEMENTARY TABLES

**Table S1.** Characteristics of LRdb versions. Note a reduction in the 2025 version due to a stricter manual filtering step.

| | # LRIs | # Ligands | # Receptors |
|---|---|---|---|
| 2019 | 3239 | 786 | 730 |
| 2024 | 3639 | 823 | 721 |
| 2025 | 3361 | 761 | 702 |

**Table S2.** Number of cells in the CD40L-CD40 dataset by Wilk, *et al*. (2024).

| Experiment | B cells | NK cells | T cells | Monocytes |
|---|---|---|---|---|
| CD40L_CD40 | 2349 | 2983 | 378 | 291 |
| CD40L_GFP | 1496 | 2577 | 295 | 277 |
| GFP_GFP | 1411 | 3087 | 333 | 187 |

**Table S3.** Lung cell population sizes

| Cell populations | Nawijn_2021 | Barbry_2020 |
|---|---|---|
| Mast cell | 595 | 158 |
| B cell | 835 | 55 |
| Mucus Secreting Cell | 55 | 272 |
| Respiratory Basal Cell | 4291 | 28027 |
| Plasma Cell | 373 | 59 |
| Multiciliate Epithelial Cell | 121 | 165 |

### SUPPLEMENTARY RESULTS AND FIGURES

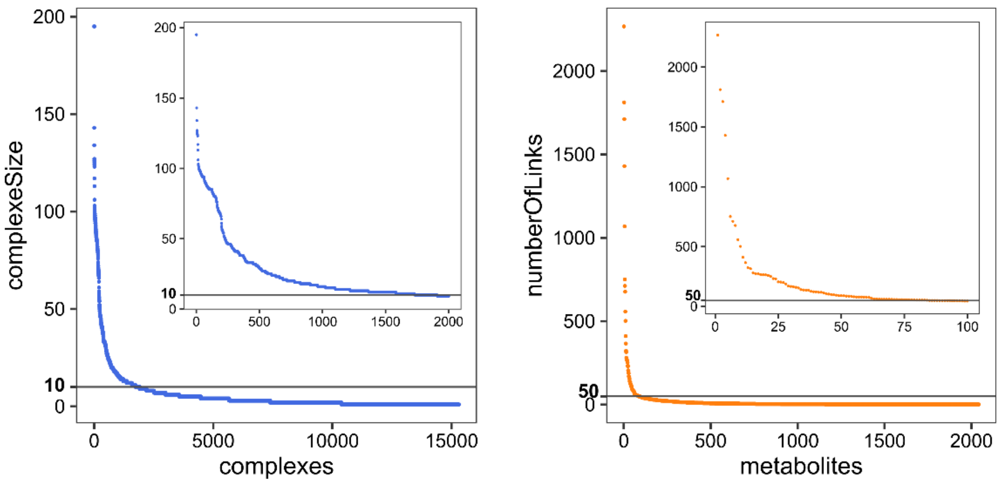


**Figure S1.** Threshold effects on the number of (**A**) protein complexes and (**B**) metabolites retained in the reference interactome. Insets = zoomed plots.

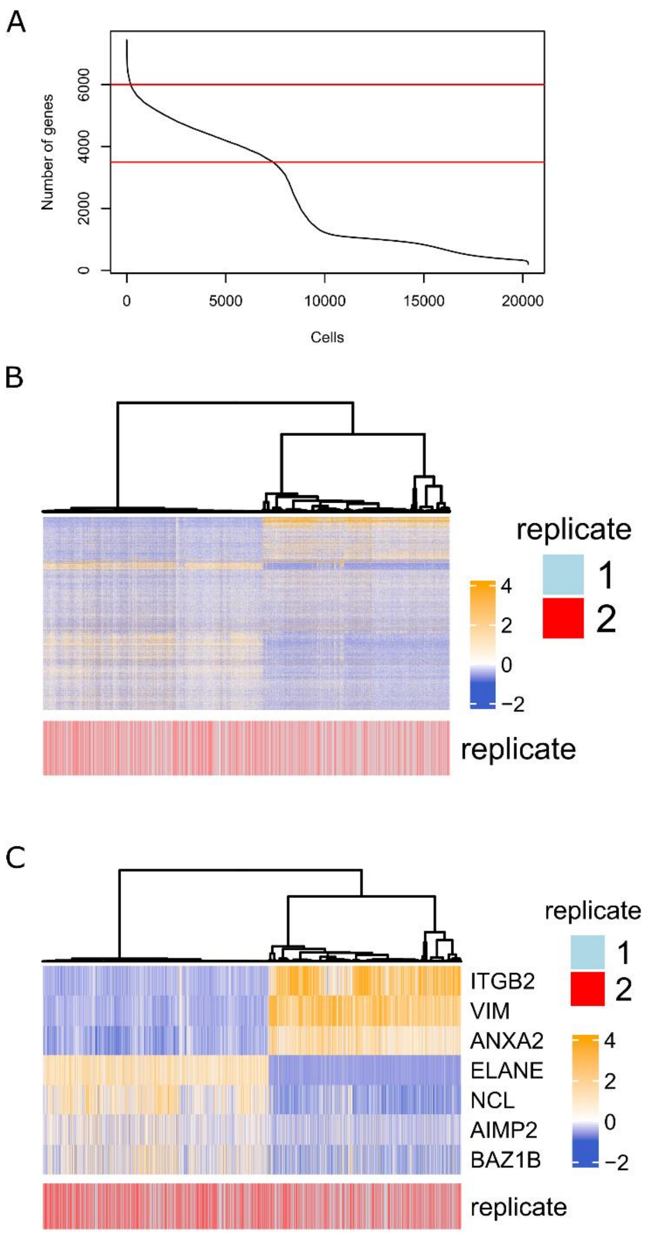


**Figure S2.** Overview of MONO-MACRO SCT data [1]. (**A**) Number of detected genes (UMI > 0) *per* cell and selection (red horizontal lines). (**B**) Gene expression overview (genes in the top 1,000 coefficient of variation). Note that SCT data were obtained from two replicates and no batch effect was observed (no batch effect correction applied). (**C**) Sorting according to the average expression of *VIM*, *ITGB2*, *NCL*, and *ELANE*.

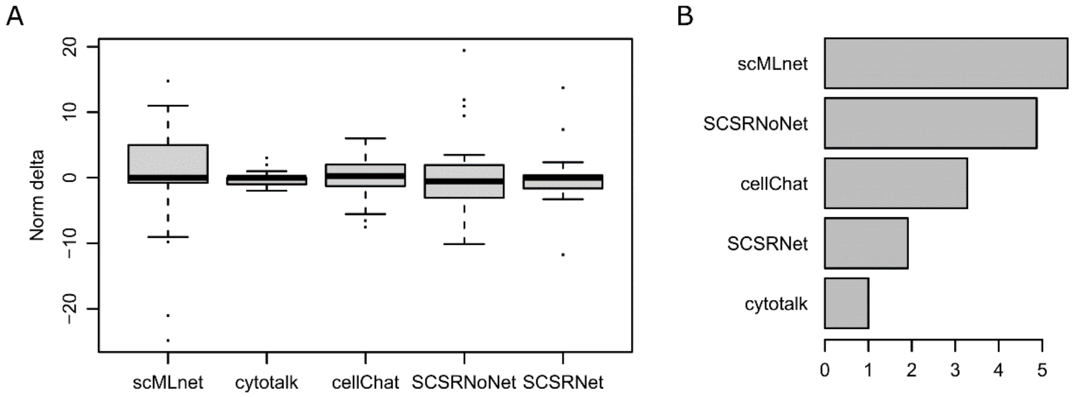


**Figure S3.** (**A**) Absolute normalized differences between numbers of paracrine LRIs identified in matching populations of the two lung atlases separately. (**B**) Corresponding interquartile differences.

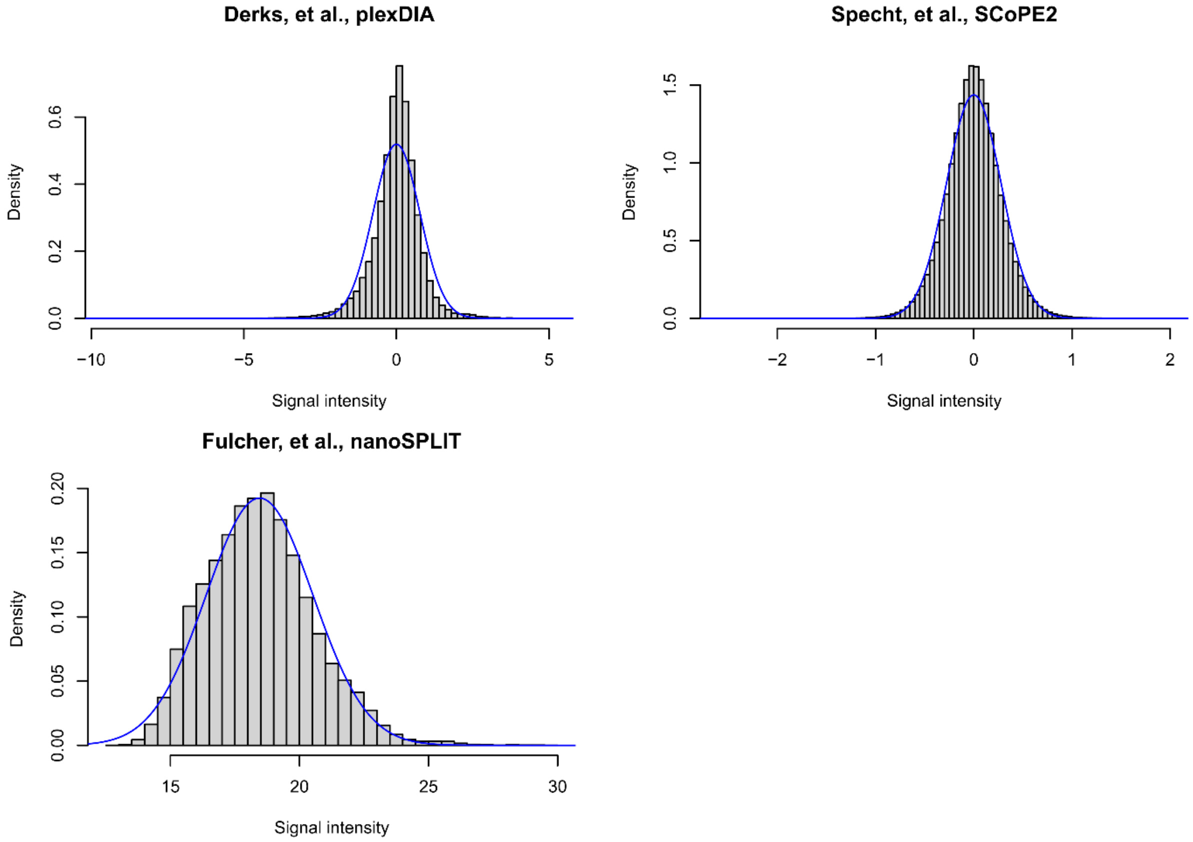


**Figure S4.** Illustrative distributions of scProt-MS protein expression values for the DDA SCoPE2 [1] and DIA plexDIA [2] workflows. The same distributions are obtained with the SCoPE-MS [3] and SCeptre [4] workflows. NanoSPLIT data led to a similar distribution but shifted in positive values far from zero [5]. Blue solid line, the closest (maximum likelihood) normal distribution for reference. As we neither assumed nor used normality in our study, this curve is only indicative and it essentially shows that the data do not distribute normally. We also used the broad A-stable family of distributions and obtained a better fit, although not accurate, indicating that scProt-MS data do not follow an A-stable distribution (fit not shown).

**ScProt-MS ligand and receptor profiles reflect cell identities**

To evaluate single-cell ligand and receptor data, we used a dataset that was obtained using OCI-AML8277 cells, an *in vitro* model of acute myeloid leukemia (AML) [4,6]. This model reproduces the hierarchy of cells that are typically observed in AML: self-renewing leukemic stem cells (LSC), progenitors, and terminally differentiated blasts. We called this dataset AML-DIFF. Using the SCeptre workflow that relies on isobaric tags (see Fig. 3A) [4], the authors obtained the individual proteomes of 255 cells that covered 1,129 proteins, among which 43 were ligands and 13 were receptors. The authors of the AML-DIFF dataset were interested in separating blasts from progenitors and LSCs. By differential analysis, they selected proteins associated with these cell types. Using their protein selection, we applied hierarchical clustering to obtain two cell clusters. The first cluster contained only blasts, except for one cell (Fig. S5A). The second cluster contained a mixture of progenitors, LSCs, and blasts with an intermediary phenotype. Then, we increased the number of clusters to improve partitioning. With four clusters, we could isolate the misclassified blasts and some LSCs and progenitors. However, this partition could not separate progenitors from LSCs, the majority of which were left in a large cluster. Conversely, by simply focusing on ligands and receptors, thus ignoring the authors' original protein selection, a two-cluster partition could separate blasts from progenitors and LSCs equally well (Fig. S5B). By increasing the number of clusters to four, we obtained one cluster dominated by LSCs and another cluster that contained mostly progenitors, but also some misclassified blasts and LSCs (Fig. S5B). Therefore, the expression of ligands and receptors was enough to provide a reasonable separation of the three cell types. To measure the agreement between the partitions obtained with the original protein selection and with only ligands and receptors, we used two standard measures of partition similarity: the Rand Index and the V-measure (or normalized mutual information). The obtained index values were similar and they were all significantly higher than random permutation-generated null distributions (Fig. S5C).

These results suggested that substantial information about cell identity was specifically carried by ligands and receptors. To confirm this hypothesis, we used quantitative proteomic data generated for 378 cell lines of the Cancer Cell Line Encyclopedia (CCLE) [7]. We checked whether the expression level of ligands and receptors was more variable across cell lines compared with that of proteins that were not secreted or not at the plasma membrane (i.e. within proteins). In this dataset, expression was given as a ratio relative to a reference mixed proteome, $\log_2$-transformed and corrected for batch effect (Fig. S5D). As the expression values were on similar scales for all proteins and the average expression was close to zero, we did not estimate variability using the coefficient of variation, but using the standard deviation directly (Fig. S5E). Ligands and receptors displayed significantly increased variability compared with within proteins. Plasma membrane and secreted proteins that were neither ligands nor receptors harbored intermediate variability. We obtained similar results also using CCLE transcriptomic data (Fig. S6).

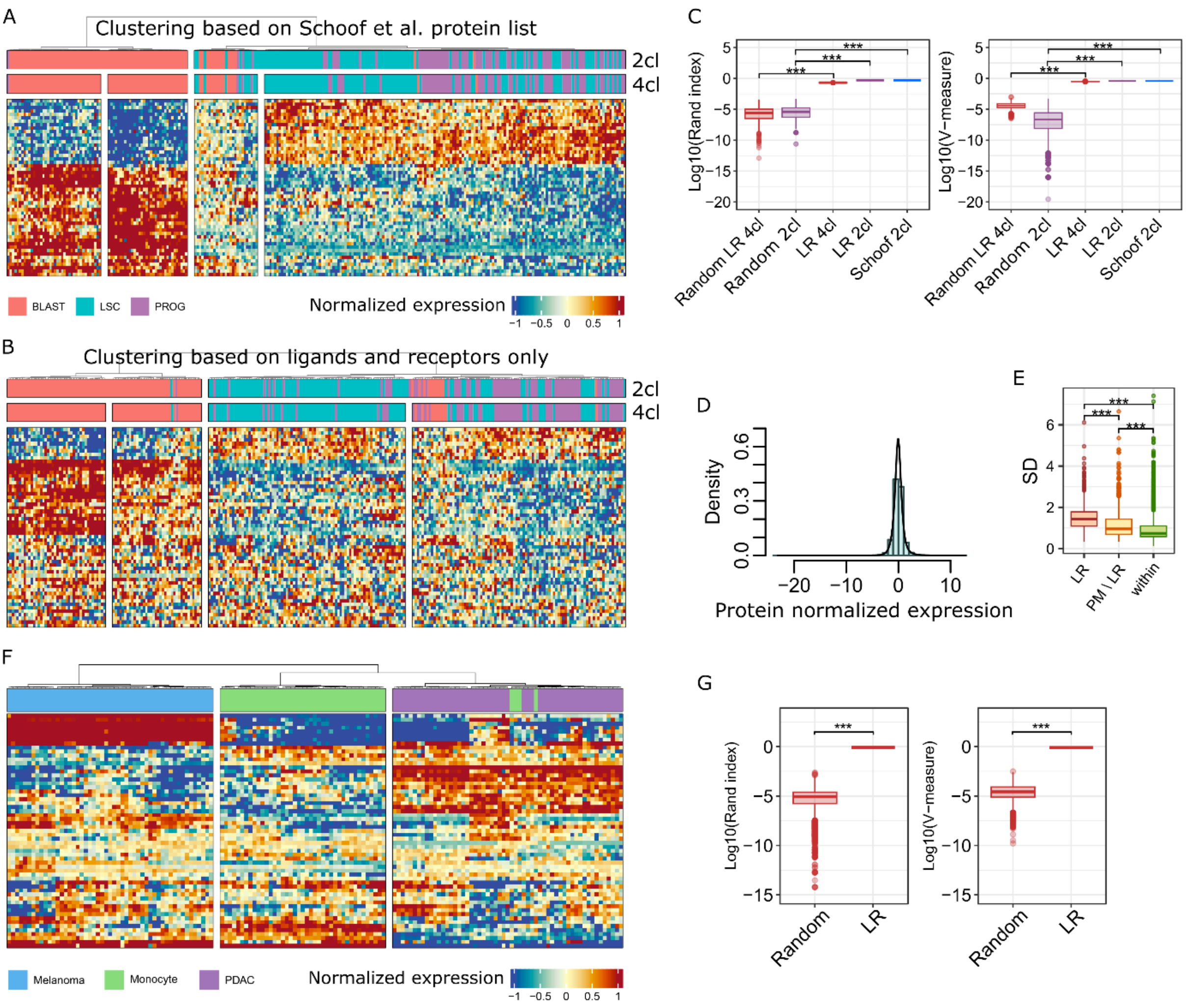


**Figure S5.** Discriminatory potential of ligands and receptors. (**A**) Cell clustering based on the proteins identified in the AML-DIFF dataset and used by the authors to distinguish blasts from leukemic stem cells (LSC) and progenitors (Prog) [4]. 2cl, two-cluster partition and 4cl, four-cluster partition. (**B**) Clustering based on all the ligands and receptors found in the AML-DIFF dataset. (**C**) Comparison of partitions in (A) and (B) to a perfect partition using the Rand Index and V-measure yielded significantly higher values than randomized partitions. (**D**) Normalized expression of proteins in 378 cancer cell lines (CCLE). (**E**) Expression variability (SD) of ligands and receptors (LR), plasma membrane proteins (GO:0005886) without LR (PM\LR), and non-plasma membrane proteins (within) in the 378 cancer cell lines. (**F**) Cell clustering (CC-MONO dataset [2]) based only on the expression levels of ligands and receptors achieved an almost perfect separation of the three cell populations: melanoma cells, monocytes, and pancreatic ductal adenocarcinoma (PDAC) cells. (**G**) Significance of the CC-MONO clustering partition. For both datasets: *** $P < 10^{-15}$ (Wilcoxon test).

We then considered a second scProt-MS dataset, which we named CC-MONO. This dataset was acquired using a mixture of pancreatic ductal adenocarcinoma cells, melanoma cells and monocytes [2]. Its depth was comparable to that of the AML-DIFF dataset, but used a (3-plexed) DIA workflow (Fig. 3A). As done for the AML-DIFF dataset, we used only ligand and receptor expression data (51 ligands, 8 receptors) to cluster cells (Fig. S5F). Except for few monocytes, the partition of cells was perfect, as indicated also by the significant partition index P-values (Fig. S5G).

With respect to the 2019 version of *LRdb*, which we used for scProt-MS analyses and contained 3,239 LRIs, the 56 proteins (ligands or receptors) in the AML-DIFF database did not form any known LRI. In the CC-MONO dataset, the 59 proteins (ligands and receptors) formed 8 putative LRI. This was obviously too little to discuss these interactions. We checked two additional murine datasets [5,8] with comparable depth of protein quantification (1,089 and 1,405 proteins, respectively) and we obtained similar numbers (22 ligands and 14 receptors, and 35 ligands and 20 receptors, respectively).

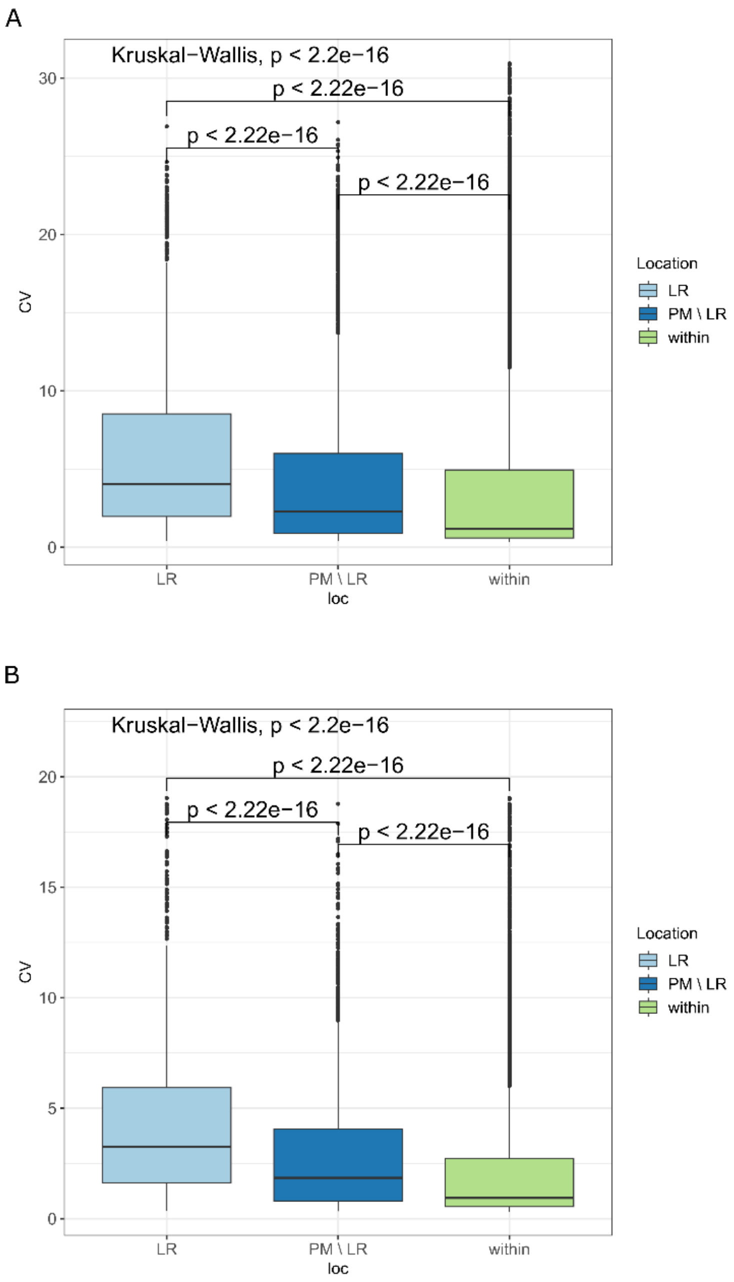


**Figure S6.** Variability of protein-coding gene expression across the different CCLE cell lines. (**A**) We used 1,021 cells lines with available RNA-seq data. The coefficient of variation (CV) reported the variability. Genes without at least a read count = 10 in 10 cell lines were discarded (n=812 genes). LR, genes that in LR*db* encode a ligand or receptor; PM\LR, genes encoding a protein included in the plasma membrane GO:0005886 term, but not a ligand or receptor; within, genes encoding a protein not included in GO:0005886. We used the Wilcoxon test for pairwise median comparisons, and the Kruskal-Wallis test for comparing all protein types. (**B**) Same analysis but restricted to the CCLE cell lines with available quantitative proteomic data from Ref [7] (n=371) to check whether there was a bias (no bias detected, very similar results).

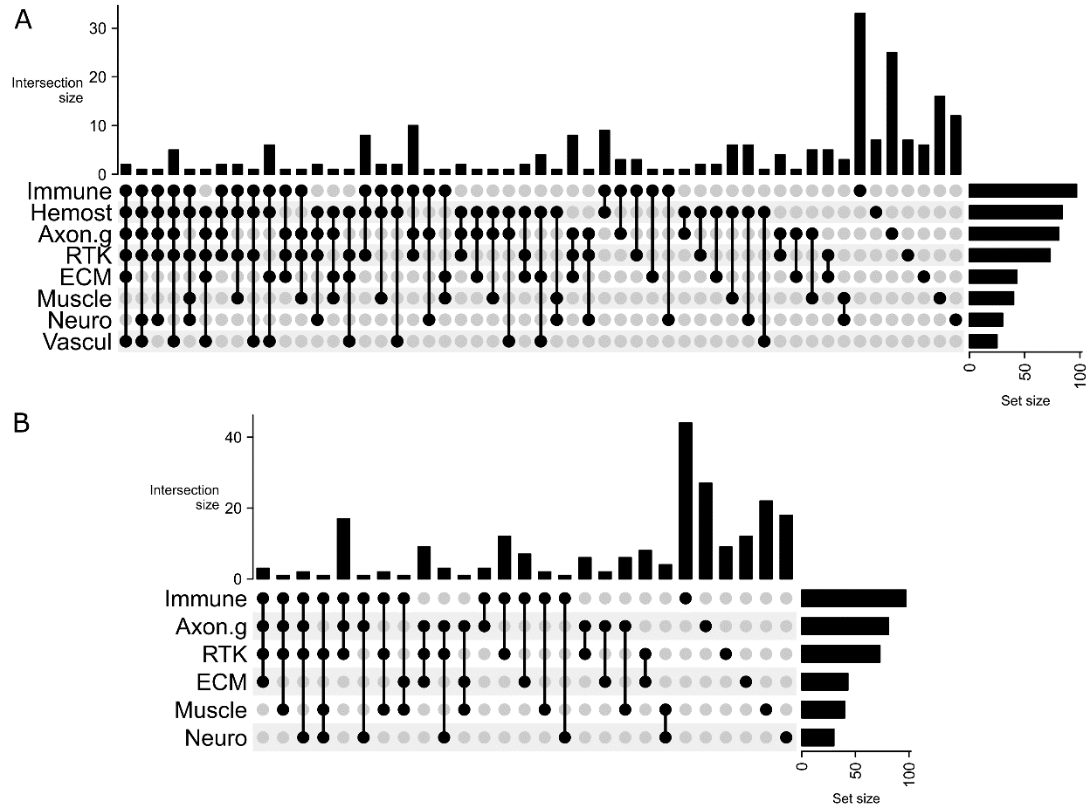


**Figure S7.** UpSet plot showing intersections between pathways in the MONO-MACRO scProt-MS dataset analyzed with SingleCellSignalR . (**A**) The eight most covered pathways. (**B**) The six pathways retained in the autocrine scProt-MS macrophage network. Immune, adaptive immune system and neutrophil degranulation pathway (Reactome IDs R-HSA-1280218 and R-HSA-6798695); Hemost, hemostasis pathway (R-HSA-109582); RTK, signaling by receptor tyrosine kinase pathway (R-HSA-9006934); ECM, extracellular matrix remodeling pathway (R-HSA-1474244, R-HSA-3000178, R-HSA-1474228, R-HSA-3000171); Muscle, muscle contraction pathway (R-HSA-397014, R-HSA-445355); Neuro, neuronal system pathway (R-HSA-112316); Vascul, cell surface interactions at the vascular wall (R-HSA-202733), Axon.g, axon guidance (R-HSA-422475).

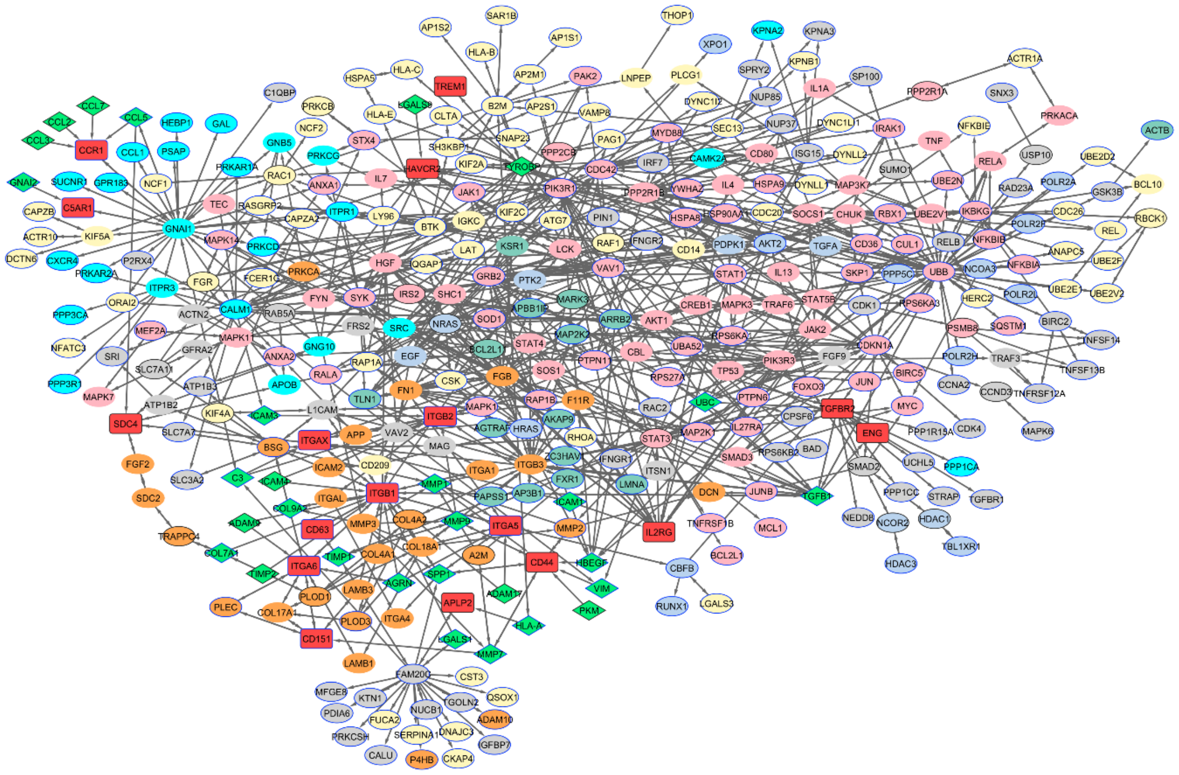


**Figure S8.** scRNA-seq macrophage autocrine network (with all the gene names).

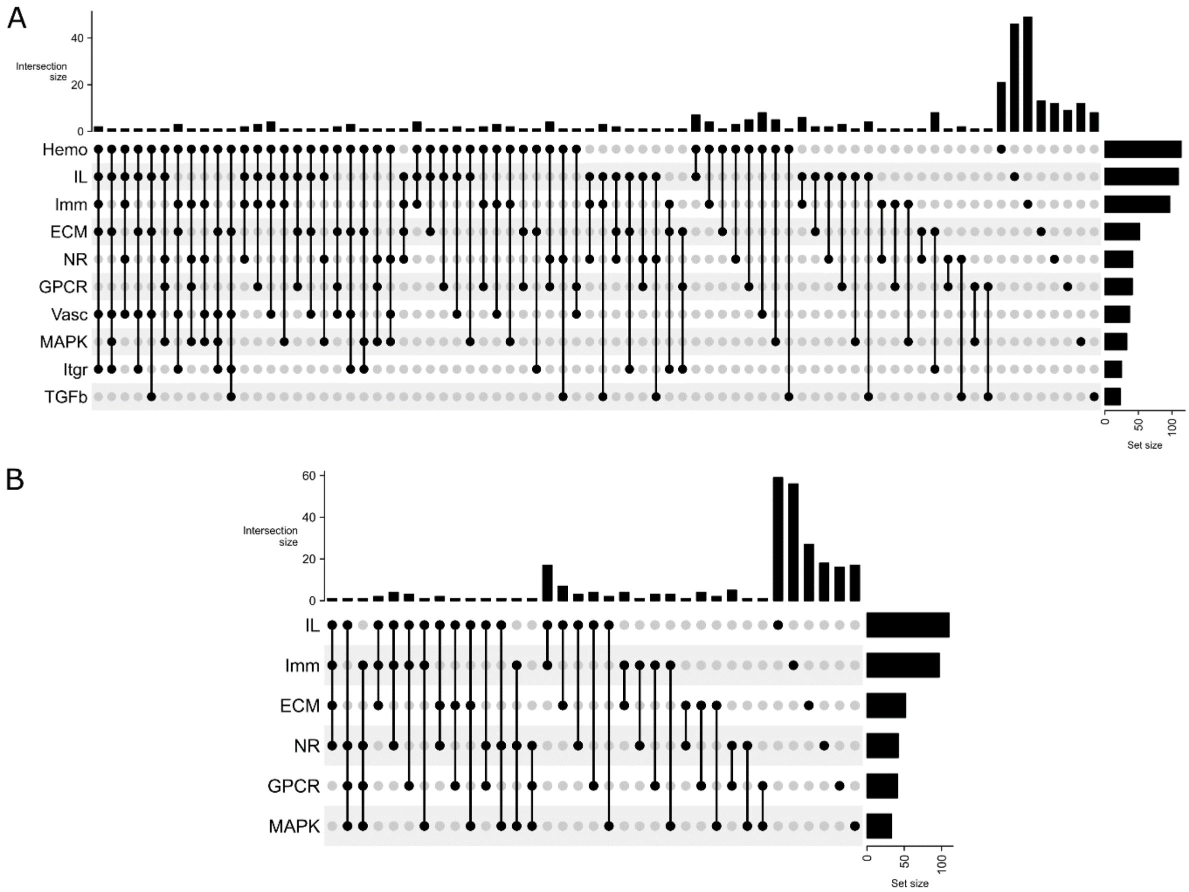


**Figure S9.** UpSet plots for MONO-MACRO transcriptomics data [1]. (**A**) All the main pathways. (**B**) Selected pathways to annotate the autocrine transcriptomic network in macrophages. Immune, adaptive immune system and neutrophil degranulation pathway (Reactome IDs R-HSA-1280218 and R-HSA-6798695); Hemo, hemostasis (R-HSA-109582); IL, Interleukin signaling (R-HSA-449147); NR, nuclear receptors (R-HSA-9006931); GPCR, G alpha (i) signalling events (R-HSA-418594); ECM, extracellular matrix remodeling (R-HSA-1474244, R-HSA-3000178, R-HSA-1474228, R-HSA-3000171); Vasc, cell surface interactions at the vascular wall (R-HSA-202733); TGFb, TGF-beta signaling (R-HSA-170834, R-HSA-9006936, R-HSA-2173789, R-HSA-2173791); ITGR, Integrin cell surface interactions (R-HSA-216083); MAPK, Oncogenic MAPK signaling (R-HSA-6802957).

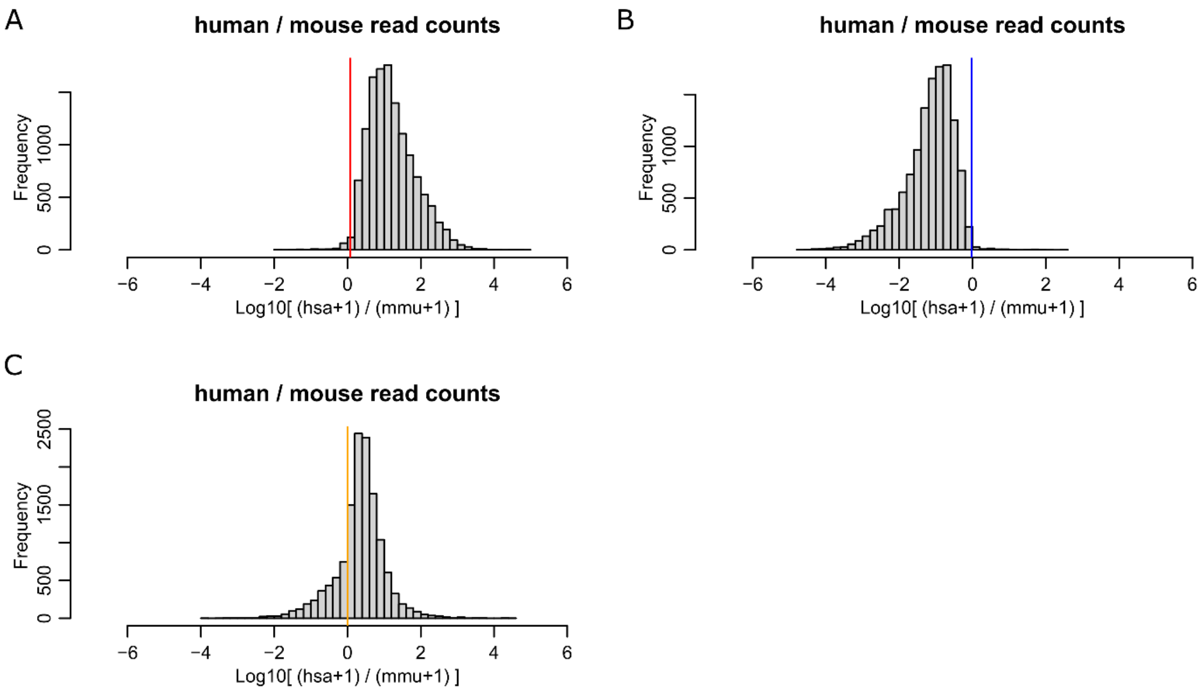


**Figure S10.** Separation of murine and human transcripts based on the log-ratio of read counts against the two genomes. Mouse genes were mapped to their human orthologs to compare human and mouse counts for a given gene. Human and mouse genes with no orthologs were discarded. (**A**) Human control sample displayed almost no match against the mouse genome. First percentile of the log-ratios was at 0.0058 (red line). (**B**) Same analysis for the mouse control sample. Ninety-ninth percentile was 0.0060 (blue line). (**C**) Same analysis for PDX samples, where a mixture of human and mouse genes are expressed. Based on (A) and (B), we set the practical limit at 0 to distinguish between mouse *versus* human cell expressed genes. Here, the average read count across all the PDX (in each species) was used for computing the log-ratios. More sophisticated methods of assigning the species exist [9], but for the purpose of this application example we preferred to stick to elementary processing.

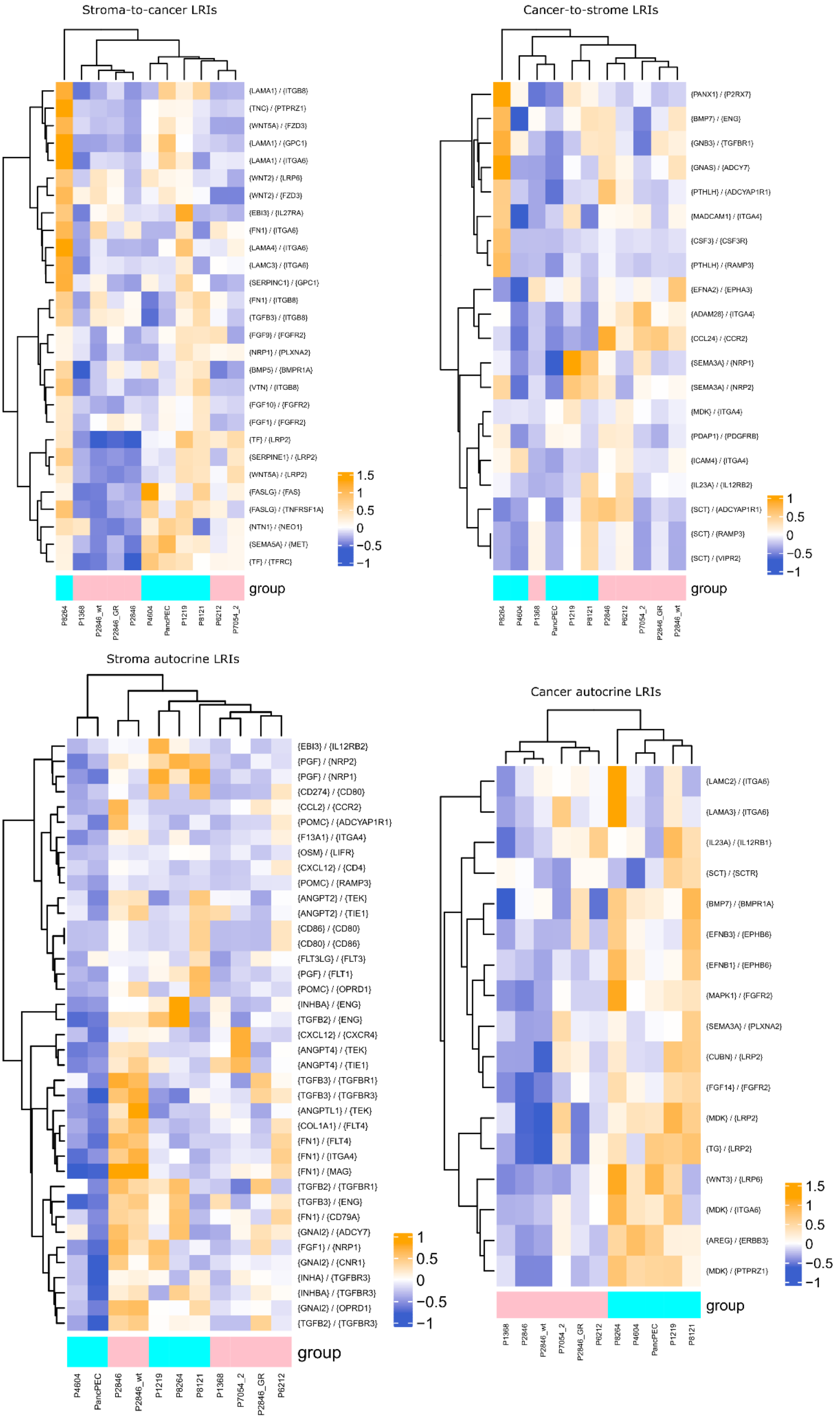


**Figure S11.** All LRIs found in the PDX data.

### SUPPLEMENTARY REFERENCES

1. Specht H, Emmott E, Petelski AA, et al. Single-cell proteomic and transcriptomic analysis of macrophage heterogeneity using SCoPE2. Genome Biol. 2021; 22:50
2. Derks J, Leduc A, Wallmann G, et al. Increasing the throughput of sensitive proteomics by plexDIA. Nat. Biotechnol. 2023; 41:50–59
3. Budnik B, Levy E, Harmange G, et al. SCoPE-MS: mass spectrometry of single mammalian cells quantifies proteome heterogeneity during cell differentiation. Genome Biol. 2018; 19:161
4. Schoof EM, Furtwängler B, Üresin N, et al. Quantitative single-cell proteomics as a tool to characterize cellular hierarchies. Nat. Commun. 2021; 12:3341
5. Fulcher JM, Markillie LM, Mitchell HD, et al. Parallel measurement of transcriptomes and proteomes from same single cells using nanodroplet splitting. 2022; 2022.05.17.492137
6. Lechman ER, Gentner B, Ng SWK, et al. miR-126 Regulates Distinct Self-Renewal Outcomes in Normal and Malignant Hematopoietic Stem Cells. Cancer Cell 2016; 29:214–228
7. Nusinow DP, Szpyt J, Ghandi M, et al. Quantitative Proteomics of the Cancer Cell Line Encyclopedia. Cell 2020; 180:387-402.e16
8. Petrosius V, Aragon-Fernandez P, Üresin N, et al. Exploration of cell state heterogeneity using single-cell proteomics through sensitivity-tailored data-independent acquisition. Nat. Commun. 2023; 14:5910
9. Kluin RJC, Kemper K, Kuilman T, et al. XenofilteR: computational deconvolution of mouse and human reads in tumor xenograft sequence data. BMC Bioinformatics 2018; 19:366
